# Food web context modifies predator foraging and weakens trophic interaction strength

**DOI:** 10.1101/2024.03.04.583297

**Authors:** Kimberley D. Lemmen, Frank Pennekamp

**Affiliations:** Department of Evolutionary Biology and Environmental Studies, University of Zurich, Zurich, Switzerland

**Keywords:** trophic interaction modification, higher-order interaction, species interactions, predator– prey model, trait mediated indirect effects, community dynamics, functional response

## Abstract

Trophic interaction modifications (TIM) are widespread in natural systems and occur when a third species indirectly alters the strength of a trophic interaction. Past studies have focused on documenting the existence and relative magnitude of TIMs, however the underlying processes and long-term consequences often remain elusive. To address this gap, we experimentally quantified the density-dependent effect of a third species on a predator’s functional response to identify the processes impacted by, and consequences of TIMs. To do so we conducted short-term experiments with two ciliate communities each composed of a predator, prey, and non-consumable ‘modifier’ species. In both communities, increasing modifier density weakened the trophic interaction strength, due to a negative effect on the predator’s search clearance rate, however the magnitude of the effect differed with prey species identity. Using parameters estimated from our experimental observation we simulated long-term dynamics and observed quantitative differences between models that account for TIMs or include only pairwise interactions. Our study is a clear demonstration that TIMs are important to understand and predict community dynamics and highlights the need to extend past pairs of focal species to understand the consequences of species interactions in communities.

## Introduction

Trophic interactions play a key role in structuring communities and determine their dynamics and stability (Terborgh 2015). Trophic interactions are commonly studied by (experimentally) decomposing a community into focal pairs that interact in a direct pairwise manner (Arditi & Ginzburg 2012). This reductionist approach assumes that the dynamics of a community can be fully explained by accounting for all the pairwise interactions within (Levine *et al*. 2017). However, this breaks down if the presence of additional species modifies how a focal species pair interacts with each other (Wootton 1993).

Community models that only account for pairwise interactions often fail to predict the quantitative dynamics observed in real-world communities (Mayfield & Stouffer 2017; Mickalide & Kuehn 2019) and produce erroneous qualitative predictions (“ecological surprises” such as extinction vs. coexistence of community members) (Fox 2023). Properly accounting for interaction modifications is therefore a prime challenge for improved predictions of the dynamics and the qualitative state of ecological communities.

Trophic interaction modifications (TIM) occur when the presence of a third species indirectly affects the direct interaction between a consumer (e.g., predator, parasitoid, detritivore) and its resource (Terry *et al*. 2017). TIMs are a category of higher-order interactions (Billick & Case 1994) and encompass all indirect effects that modify consumption and may manifest through a variety of ecological and evolutionary processes (Barbosa *et al*. 2009; Holland *et al*. 2005). One of the most studied classes of TIMs is the effect of fear (Werner & Peacor 2003; Schmitz *et al*. 2004; Wirsing *et al*. 2021), in which an intermediatory consumer is vulnerable to the top predator when foraging. When the consumer detects the presence of a top predator it reduces its foraging effort to decrease its vulnerability, this behavioural change also decreases its consumption. Thus, the top predator can influence the strength of the consumer-resource interaction outside of a direct consumptive effect (Schmitz *et al*. 1997; Preisser *et al*. 2005; Werner & Peacor 2006). A TIM that weakens consumer-resource interactions will stabilize community dynamics and allow for the persistence of the resource species in the presence of a third species (Vos *et al*. 2001; van Veen *et al*. 2005; Hammill *et al*. 2015).

Empirical studies have provided evidence for TIMs in natural systems (Werner & Peacor 2003; Kratina *et al*. 2007; Kehoe *et al*. 2016; Wasserman *et al*. 2016). These studies often focus on short-term effects, for example, if changes in predator behaviour are due to modifications in resource traits induced by a third species (i.e., trait-mediated indirect effects, (Schoener & Spiller 2012). However, such studies often cannot inform about long-term community consequences such as the persistence of a focal consumer-resource pair (Abrams 1995) hampering our empirical understanding regarding the effects of TIMs on community predictability and stability. Understanding the dynamic consequences of TIMs requires quantifying the strength of the modification in a manner that can be used in community dynamic models (Okuyama & Bolker 2007). In the past, studies often used a simple factorial design to measure changes in resource species abundance in the presence and absence of a modifier. While this information is crucial to documenting the presence of a TIM, it cannot be used to parameterize a dynamic model to predict community dynamics, which are an important requisite to support decision making (Paine *et al*. 2018).

Terry *et al*. (2017) proposed a framework to rigorously detect TIMs and understand their dynamic consequences. They argued that response surfaces across density gradients of the involved species are needed to quantitatively describe TIMs and predict their long-term effects. Previously, such experiments were prohibitively logistically complex, however rapid technological and analytical advancements have allowed for large-scale cross-gradient designs to become feasible (Pennekamp & Schtickzelle 2013; Rosenbaum & Rall 2018). An additional challenge in understanding TIMs is that the existing models used to empirically quantify their effect and strength are purely phenomenological (Letten & Stouffer 2019).

We thus have limited information on the processes underlying the interaction modification, which constrains the model’s predictive power when extrapolating beyond previously observed conditions (Bolker 2008). Terry *et al*.’s (2017) framework suggests fitting a suite of models with different formulations encoding contrasting hypotheses about the processes underlying the TIM to the response surface data. By comparing how well different formulations describe the collected data one can gain insights into biological processes driving the species interactions. This framework is a useful starting point to empirically document their presence, strength, and mechanistic origin in a manner that can be used to model community dynamics and advance theoretical efforts.

The functional response is a key ingredient of predator-prey models and describes the relationship between predator foraging rate and prey abundance (DeLong 2021; Abrams 2022). The strength of a trophic interaction is broadly determined by two processes (i) the rate at which a predator clears a given space of prey and (ii) how long it takes to process the prey (Box 1). Fundamentally, the functional response is a pairwise model (even when used to model a generalist predator), and there is limited insight into how functional response processes are affected by a modifier species (Figure 1b). Given the widespread use of functional responses to quantify trophic interactions, we need to take the first steps in understanding how they are impacted by modifier species.

**Figure 1:**
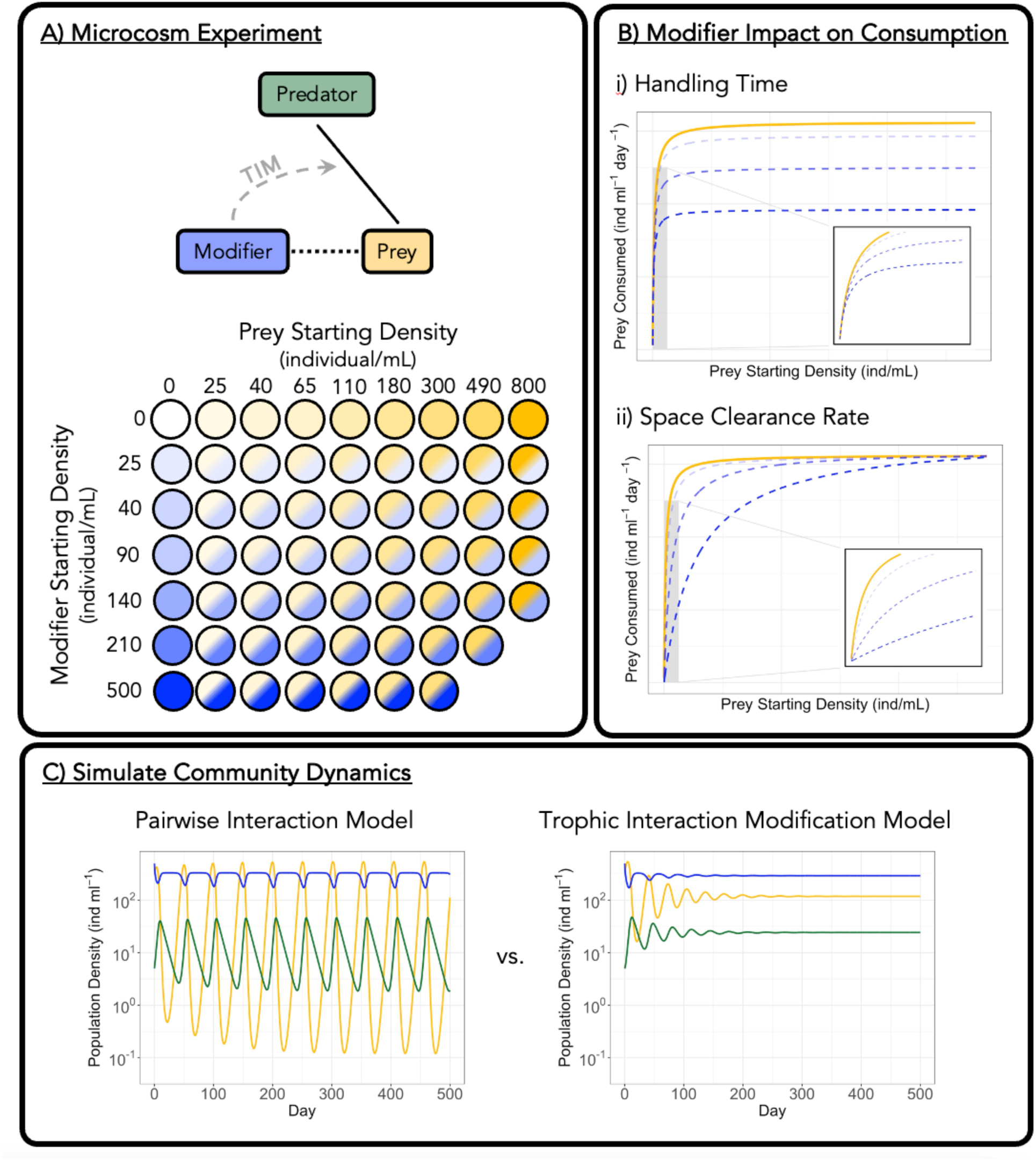
(A) Experimental approach used in this study to characterize trophic interaction modifications. In the community composition schematic black solid lines represent a trophic interaction, black short-dashed lines represent a competitive interaction and grey long-dashed dash lines represent trophic interaction modification. In the schematic of the experimental design, darker colours represent higher density of the species. (B) Visualization of the effect of trophic interaction modifications on either the space clearance rate or handling time on a predator’s functional response. Note that modifications of the space clearance rate have a large effect at low prey densities and change the rate at which the curve approaches saturation. In contrast, modifications in handling time have small effects at low prey densities, but changes the density at which the curve saturates. For simplicity both trophic interaction modifications visualized decrease consumption, however the modifier may also increase consumption. (C) Example of community dynamics of a three species system containing a predator (green), prey (yellow) and modifier (blue) speciess, we tested the Terry *et al*. (2017) framework with ciliate communities containing a predator (*Spathidium* sp.), a prey (either *Colpidium striatum* or *Dexiostoma campylum*), and a non-consumable modifier species (*Paramecium caudatum*) (Figure 1a). For each community, we combined the modifier and prey species at various densities in a factorial manner (Figure 1a) and measured the change in density over a 24-hour window with and without the predator. From these observations, we were able to construct a response surface of changes in density due to intra- and inter-specific competition and consumption. We fit different models to the response surface that, building up from a simple Type II functional response, incorporated terms that accounted for a modifier-dependent change in the space clearance rate, handling time, and predator’s time budget (Figure 1b). The parameter estimates from the best model of each community were then used to simulate its dynamics over an extended time frame (Figure 1c).

We predict that the presence of the modifier will result in a density-dependent decrease in the space clearance rate and an increase in the handling time of the predator. The combined effect of which will significantly reduce consumption at high modifier densities. We further predict that the magnitude of the modifier effect will differ between species, and will be greater for *Colpidium* due to its phenotypic similarities (i.e., size) with the modifier. Finally, we predict that the dynamics of the community will display increased stability and predictability when we account for the effect of the modifier than when pairwise interactions alone are used to model the system.

## Methods

### Data Collection – functional response experiment

To examine the role of TIMs, we constructed experimental ciliate communities consisting of a predator, prey, modifier species, and a bacteria basal resource (Fig 1a). In this study, the modifier species cannot be consumed by the predator, and it competes with the bacterivorous prey species for the basal resource. To look for generalities in the effect of TIMs we used two different community compositions. In both communities we used *Spathidium* sp. (40 - 300 um) as the predator, *Paramecium caudatum* (170–300 um) as the modifier and a mix of three bacteria species (*Bacillus subtilis*, *Serratia fonticola*, and *Brevibacillus brevis*) as the basal resource. The communities differed in prey species identity, *Colpidium striatum* (50-100 um) and *Dexiostoma campylum* (35-90 um). All ciliate species (hereafter referred to by their genus name) have been previously used to investigate trophic interactions (e.g., Daugaard *et al*. 2019; Shen *et al*. 2023). Protists were cultured to standardize the population state before the experimental trials (Appendix S1a).

To characterize the effect of the modifier on predator consumption, we incubated experimental units containing the prey, modifier, or both species, with and without the predator. For the functional response trials, we combined different starting densities of the prey (nine densities) and modifier (seven densities) in a factorial manner with a density of five predators per mL (Figure 1a). All trials were conducted in 1mL of media on 24-well plates. Each combination was replicated six times (high prey density combinations of both species were not logistically possible, Appendix S1b) and randomly positioned on the plate. To account for changes in both prey and modifier population density due to growth, intra- and interspecific competition, we incubated experimental units of all prey-modifier densities combinations without predators, each combination was replicated three times. To initiate a trial, prey and modifier densities were manipulated via the dilution of the stock culture, and predators were manually pipetted from the holding plate. The presence of the five predators was visually confirmed before adding the other species. As all experimental units could not be conducted on a single day (*Colpidium* community: 534 units, *Dexiostoma* community: 518 units), we used a block design over six days for each community. We divided the units so that the entire range of prey-modifier combinations was included in each block and that the replicates were spread evenly (Appendix S1b). Within each block, experimental units were randomly allocated to a plate and a well. Plates were incubated in the dark at 15°C for 24 hours with a jar of deionized water to minimize evaporation.

To quantify the change in population densities of the three ciliate species during the experiment we measured the density of all species at the end of the experiment. To estimate final prey and modifier abundance we took three videos (Leica M205C, 8X magnification) of a well-homogenized 0.486 mL subsample of each experimental unit (Appendix S1c). We processed the videos using the R package BEMOVI (Pennekamp *et al*. 2015). For species classification, we trained a support vector machine classifier from the R package e1071 (Meyer *et al*. 2022) with monoculture videos (Appendix S1c, Pennekamp *et al*. 2017). Final predator abundance was assessed manually using a light stereoscope (Nikon SMZ1500, 8X magnification) with 0.32mL of the sample. As all experimental units could not be processed simultaneously, we recorded the total incubation time for each unit and controlled for differences in the analysis. Units were incubated for an average of 25.7 hours (min = 24 hours, max = 27.4 hours).

### Modelling observed changes in population density

Ciliates have rapid generation times, as such, over the 24-hour experiment changes in population density of the prey species may be attributed to (i) reproduction, (ii) consumption by the predator, and (iii) competition with the modifier. To account for these dynamics, we used a system of ordinary differential equations (ODE) to describe observed changes in the density.

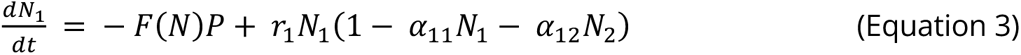

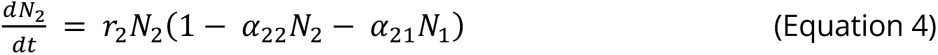

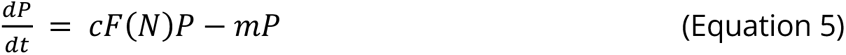

Where N_1_, N_2_ and P are the density of the prey, modifier, and predator species respectively. F(N) is the per capita consumption rate of the predator, this component of the ODE differed between model fittings (described below) allowing us to identify the general processes by which the modifier species altered the trophic interaction. The prey and modifier parameters are the growth rate (r_1_ and r_2_), intraspecific competition coeffcients (*α*_11_ and *α*2_2_) and interspecific competition coeffcients (*α*_12_ and *α*_21_) respectively. These parameter values are fixed based on values estimated from a model fit to the competition only trials (Appendix Table S2-1). The predator parameters are the efficiency of conversion of prey into predator individuals (c) and the mortality rate (m). We fixed the values of these parameters based on pre-experimental trials (Appendix S1d). By fixing all non-F(N) parameters we ensured differences between ODE fits were due solely to differences in the trophic interaction models. We tested the effect of freely estimating all parameters using the *Colpidium* community and observed quantitatively similar result.

To characterise the effect of the modifier on the trophic interaction [i.e., F(N)] we used different functional response formulations to represent predator consumption (Box 1), each with a different biological interpretation. To identify the process altered by the modifier species we extended a basic Type II functional response (Holling 1959, equation 1) to include terms that allowed modifier density to either increase or decrease predator consumption (Table 1). We hypothesize that a modifier species can impact the functional response several different ways. Modifier density may alter the predator’s space clearance rate or handling time, which could either increase or decrease consumption (Figure 1b). To model density-dependent alteration of these parameters we include a linear scaling term (Hauzy *et al*. 2010; Mocq *et al*. 2021). We acknowledge the effect of modifier density may be non-linear (i.e., saturating, Kéfi *et al*. 2012), however, we only consider linear modifications to prevent over-parameterization and potential overfitting (Burnham & Anderson 2002). Alternatively, the modifier may interact with the predator and decrease the time available to search for or handle prey, similar to predator interference (Beddington 1975; DeAngelis *et al*. 1975; Crowley & Martin 1989). To account for this modifier density-dependent loss of time we used a model (Equation 4) proposed by van Veen *et al*. (2005) that includes loss of searching time and a modification of the Crowley-Martin (Crowley & Martin 1989) model which includes loss of both searching and handling time.

**Table 1:** Functional response forms used to model the per capita rate of prey consumption by the predator. The model parameters are: prey density (N_1_), modifier density (N_2_), space clearance rate (a), linear modification of the space clearance rate (a_12_), handling time (h), linear modification of handling time (h_12_), the time wasted by the predator encountering the modifier (μ) and the prey density where the realized space clearance rate reaches half of the maximum (k_R_). TIM stands for trophic interaction modification and SCR stands for space clearance rate.

| Model | TIM<br>on a | TIM<br>on h | Density-dependent<br>SCR | Functional Response form |
| --- | --- | --- | --- | --- |
| 1. Type II | No | No | No | $\frac{aN_1}{1 + ahN_1}$ |
| 2. Type II + linear a TIM | Yes | No | No | $\frac{(a + a_{12}N_2)N_1}{1 + (a + a_{12}N_2)hN_1}$ |
| 3. Type II + linear h TIM | No | Yes | No | $\frac{aN_1}{1 + a(h + h_{12}N_2)N_1}$ |
| 4. Type II + linear a TIM + linear h TIM | Yes | Yes | No | $\frac{(a + a_{12}N_2)N_1}{1 + (a + a_{12}N_2)(h + h_{12}N_2)N_1}$ |
| 5. Type II + van Veen | Yes | No | No | $\frac{aN_1}{1 + ahN_1 + \omega N_2}$ |
| 6. Type II + van Veen + linear h TIM | Yes | Yes | No | $\frac{aN_1}{1 + a(h + h_{12}N_2)N_1 + \omega N_2}$ |
| 7. Type II + Crowley-Martin | Yes | Yes | No | $\frac{aN_1}{(1 + ahN_1)(1 + \omega N_2)}$ |
| 8. Type III | No | No | Yes | $\frac{\left(\frac{aN_1}{k_R + N_1}\right)N_1}{1 + \left(\frac{aN_1}{k_R + N_1}\right)hN_1}$ |
| 9. Type III + linear a TIM | Yes | No | Yes | $\frac{\left(\frac{(a + a_{12}N_2)N_1}{k_R + N_1}\right)N_1}{1 + \left(\frac{(a + a_{12}N_2)N_1}{k_R + N_1}\right)hN_1}$ |
| 10. Type III + linear h TIM | No | Yes | Yes | $\frac{\left(\frac{aN_1}{k_R + N_1}\right)N_1}{1 + \left(\frac{aN_1}{k_R + N_1}\right)(h + h_{12}N_2)N_1}$ |
| 11. Type III + linear a TIM + linear h TIM | Yes | Yes | Yes | $\frac{\left(\frac{(a + a_{12}N_2)N_1}{k_R + N_1}\right)N_1}{1 + \left(\frac{(a + a_{12}N_2)N_1}{k_R + N_1}\right)(h + h_{12}N_2)N_1}$ |
| 12. Type III + van Veen a TIM | Yes | No | Yes | $\frac{\left(\frac{aN_1}{k_R + N_1}\right)N_1}{1 + \left(\frac{aN_1}{k_R + N_1}\right)hN_1 + \omega N_2}$ |
| 13. Type III + Crowley-Martin | Yes | Yes | Yes | $\frac{\left(\frac{aN_1}{k_R + N_1}\right)N_1}{\left(1 + \left(\frac{aN_1}{k_R + N_1}\right)hN_1\right)(1 + \omega N_2)}$ |

Our goal was to determine if a TIM occurred in our experimental communities, and if so, which functional response formulation best described the observed change in prey density. For each hypothesized formulation of the functional response (Table 1) we fit the system of ODEs (equation 3-5) using the specific F(N) to find the parameter values that best fit the observed density change. We used a response surface approach (Okuyama & Bolker 2012) and fit the ODEs to the observations across all prey, modifier, and predation levels simultaneously (i.e., sequential fitting; Uszko *et al*. 2020). To estimate the parameter values we conducted numerical simulations and maximum likelihood estimation as described in Rosenbaum & Rall (2018) (Appendix S1d). We used the R package odeintr (Keitt, 2017) for the numerical solution of the ODEs and the bblme package (Bolker & Team, 2022) for the maximum likelihood estimation. To increase our confidence that the parameter estimates represented a true global minimum we used many different starting parameters combinations to initiate the numerical simulation (Appendix S1d). We then simulated the 95% confidence intervals for each functional response using population prediction intervals (Bolker 2008). Finally, we compared the performance of the models with contrasting functional response forms using Bayesian information criteria (Schwarz 1978). We use BIC as we are testing a defined hypothesis on a small number of models and we use data collected from a controlled experiment with well-understood predictors (Aho *et al*. 2014). This process was independently conducted for both experimental communities.

### Community Dynamics Simulation

To understand the effect on community dynamics of including TIMs we simulated the long-term population dynamics of our experimental communities using two models, one including TIMs (Table 1, models 2-7 and 9-13) and a second with only pairwise interactions (Table 1, models 1 and 8). For this comparison, we choose the model that best fit our experimental observations and the best-performing pairwise interaction model. Using the parameter values estimated by the two-model fits we simulated the community dynamics for 1100 days using equations 3-5. For each model, we estimated the 95% confidence intervals using population prediction intervals (Bolker 2008). To understand the TIM effect on long-term system dynamics, we compared measures of stability of the TIM and pairwise-only simulated communities. We assessed community stability with multiple metrics (i.e., time to extinction, amplitude and length of population oscillation, Appendix S1e). We fixed the value of c (0.007) in the simulated community dynamics. This value is larger than we used for ODE fitting. However, predator conversion effciency is diffcult to quantify over short timeframes and the value used in the simulation represents a biologically realistic value (*Spathidium*-*Dexistoma* c = 0.004-0.016 (Daugaard *et al*. 2019); various protist predator-prey c = 0-3.36 (DeLong & Vasseur 2012)). We investigated the influence of the conversion effciency on the difference between the simulated TIM and pairwise dynamics using sensitivity analysis (Appendix S1f).

## Results

### Trophic Interaction Modification Identification and Characterization

In the *Colpidium* community, modifier presence had a strong negative effect on predator consumption (Figure 2a,b). The effect of the modifier is first evident at 90 individuals/mL, negatively impacting consumption at low and intermediate prey densities. As modifier density increased so did the impacted prey range, and at the highest modifier density, the negative effect on space clearance rate almost entirely prevents consumption. For this community, the type II model which included a density-dependent effect of the modifier on the space clearance rate was the best-performing model (Table 2). Furthermore, the pairwise interaction only models were the worst-performing models.

**Figure 2:**
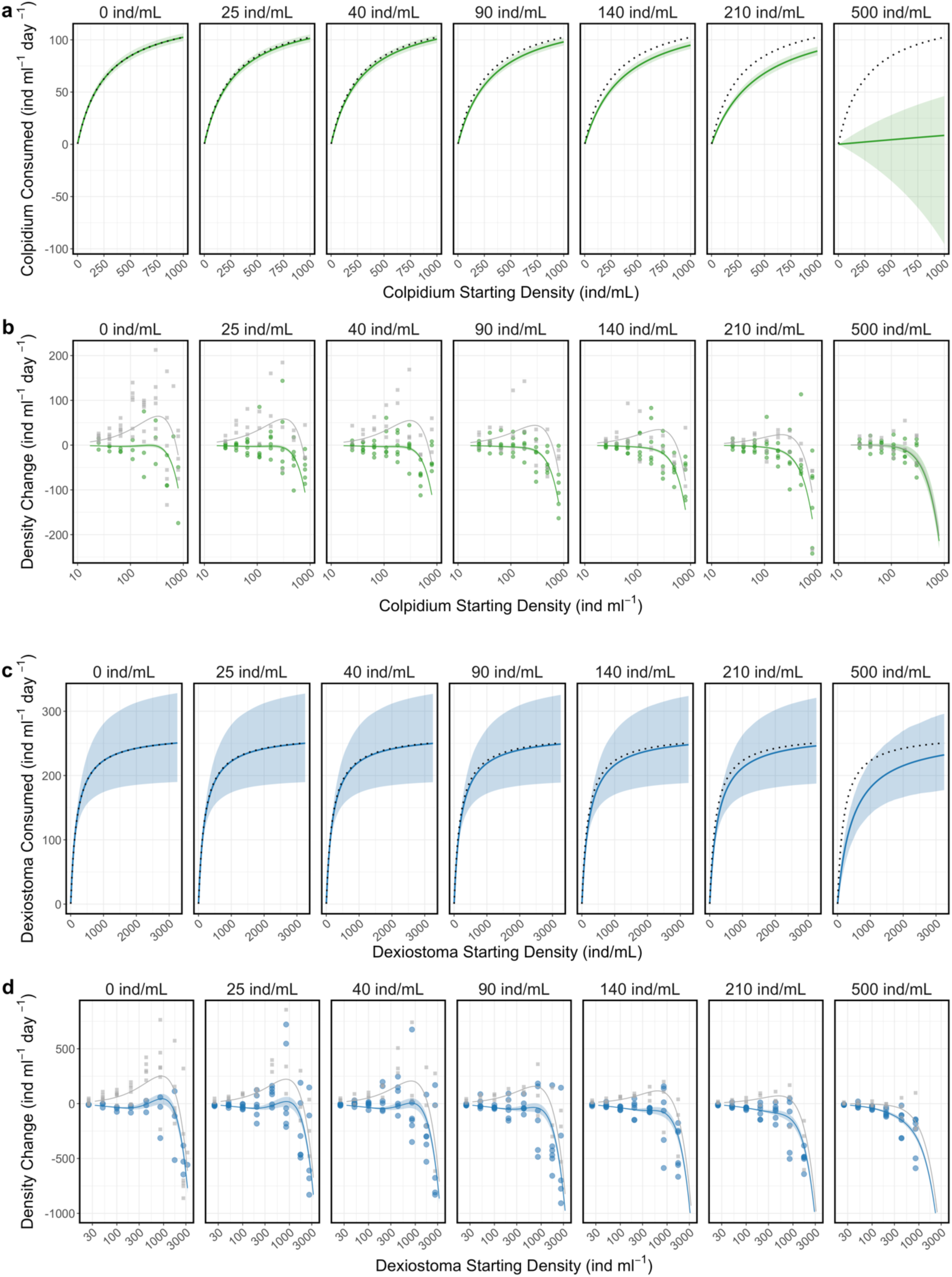
Effect of modifier density (panels) on predation (a, c) and observed change in prey density (b,d) in the *Colpidium* (a,b) and *Dexiostoma* (c,d) community. Functional response graphs visualize the number of prey predicted by the model to be consumed by predators in each community (the dotted line represents model fit with no modifier). The density changes are the observed differences in prey density over the 24-hour experiments due to predation, prey growth or intra/interspecific competition (circles are observations with predators, squares are with no predators). Observed changes in prey density may be negative and denote a reduction in prey density (i.e., consumption) or positive and denote an increase (i.e., growth). Coloured lines are the model fit with predation, and solid grey lines represent the model fit with no predation. Shaded areas represent the 95% confidence intervals of model fit (all lotka-volterra parameter values are fixed in the model as such there are no confidence intervals for the no predation model fits). Please note that the X-axis of b and d are on the log scale.

**Table 2:**
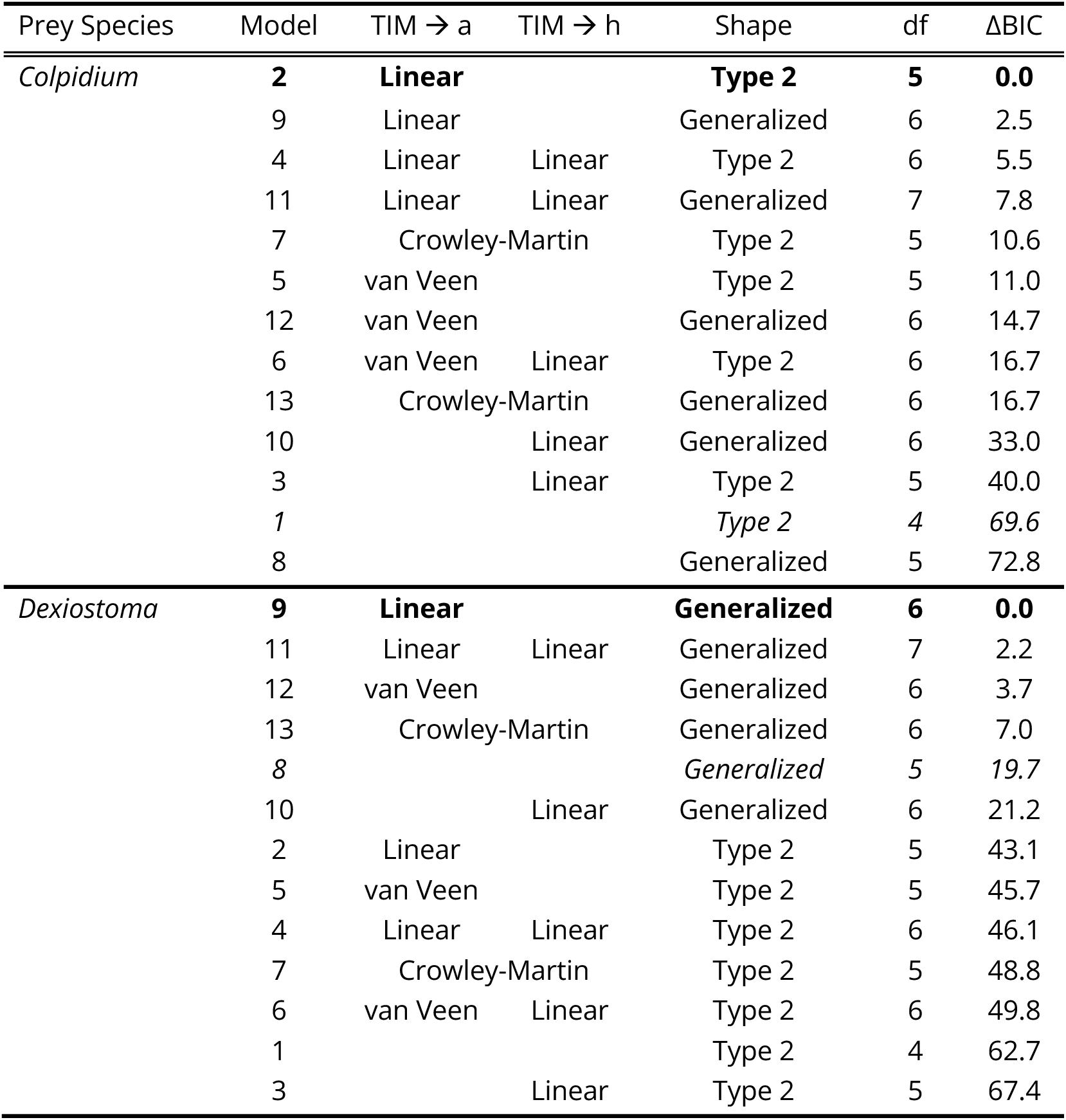
Bayesian information criterion (BIC) for the *Colpidium*-*Paramecium*-*Spathidium* and *Dexiostoma-Paramecium*-*Spathidium* communities. Functional responses considered may include a modifier effect on space clearance rate (TIM->a) and/or handling time (TIM->h) and modifier inference with the predator (van Veen and Crowley-Martin). df = degrees of freedom of the model. Models are presented within a community from lowest to highest BIC. Bold text indicates the model with the lowest BIC. Italicized text indicated the pairwise model with the lowest BIC. For estimates of the parameter values see Table S2-2.

In the *Dexiostoma* community, the modifier had a weaker effect on the predator-prey interaction (Figure 2c,d). A modifier density greater than 90 individuals per mL suppressed predator consumption at low prey densities however, this modifier effect was counteracted by high prey densities leading to almost no effect of the modifier on predator consumption when prey densities were >1000 individuals/mL. The generalized model which includes a density-dependent effect of the modifier on the space clearance rate was the best-performing model (Table 2), outperforming type II models. The model that only contained pairwise interactions was the worst-performing generalized model and the second worst type II model.

### Effect of Trophic Interaction Modification on Simulated Population Dynamics

In both communities, the simulated dynamics differed between the overall best and pairwise models. In the *Colpidium* community, the prey species is competitively excluded by the modifier, *Paramecium*, for both the TIM and pairwise model (Figure 3a). However, the time to extinction of the prey and predator significantly differed between models. In the model that accounted for a TIM, the predator went extinct earlier and the prey went extinct later than in the pairwise interaction model (Figure 3b). The sensitivity analysis suggests that higher predator conversion efficiencies resulted in greater differences in the time to extinction between pairwise and the TIM model (Appendix S3).

**Figure 3:**
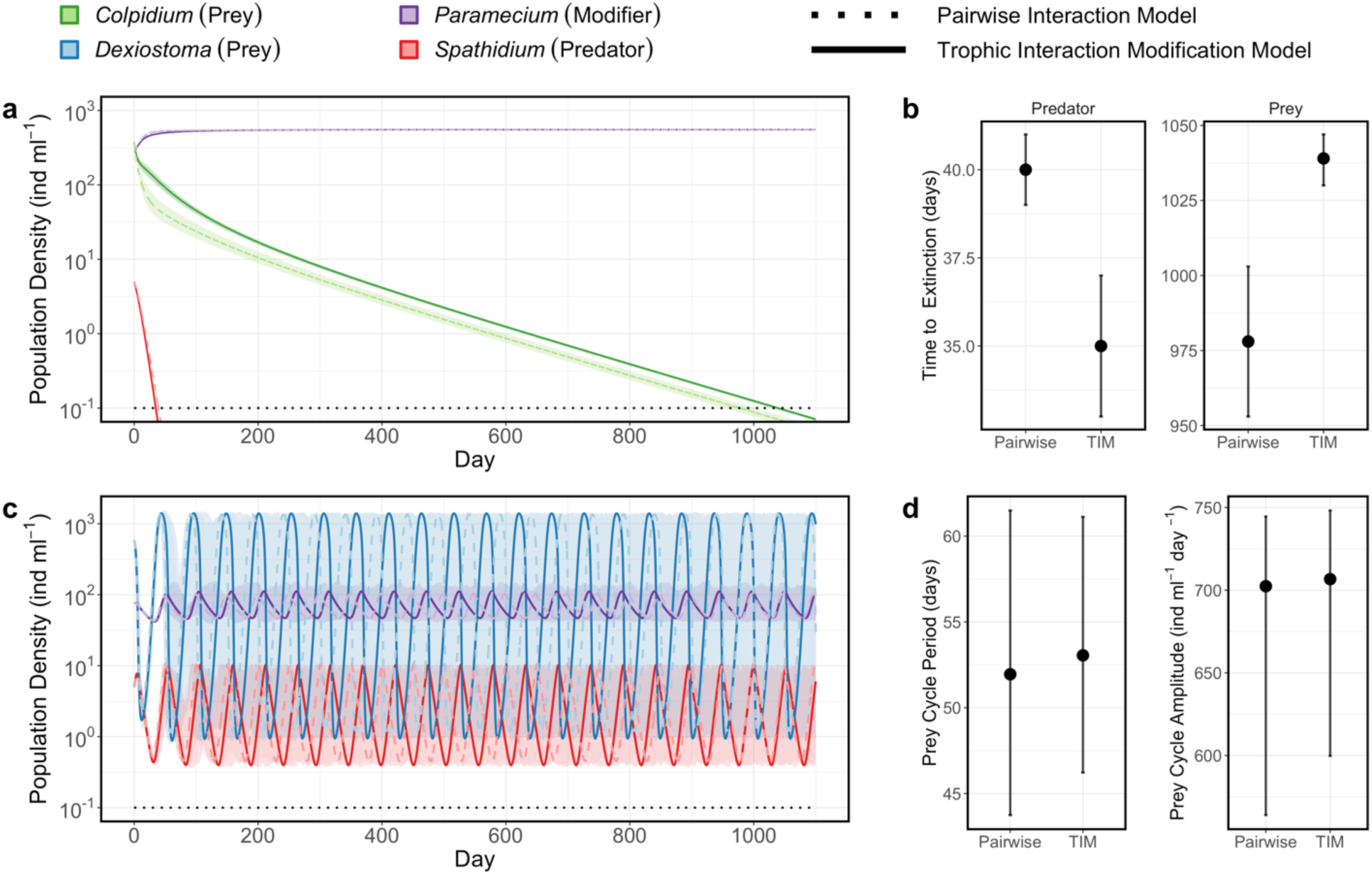
Predicted effect of trophic interaction modification on community dynamics in experimental communities (a,c) Simulated community dynamics over 1100 days based on the best model including a trophic interaction modification (TIM, solid line) and best pairwise model (dashed line). Shaded areas denote the upper and lower trend of the simulations, while the lines represent the population dynamics with the estimated parameter values. (b) The time to extinction of the predator and prey populations in the *Colpidium* community (note the difference in range of y-axis between species), error bars represent 95% confidence intervals. (d) Length and amplitude of prey cycles in the *Dexiostoma* community, error bars represent 95% confidence intervals of the cycles of the best-fit model. For both communities, the conversion rate effciency of the predator is fixed at 0.7%

In the simulated dynamics of the *Dexistoma* community, we observe stable oscillatory dynamics between the prey, modifier, and predator for both the TIM and pairwise models (Figure 3b). We observed no significant difference in the amplitude of the cycles for any of the species in the community (Figure 3d), however, the period of the cycles is longer for the TIM model. The sensitivity analysis suggests that the difference in cycle length is greatest at relatively low conversion rate effciencies.

## Discussion

Our study is the first empirical demonstration of how response surfaces in combination with a hypothesis-driven model selection approach can pinpoint the processes impacted by, and consequences of, modifier species on trophic interactions. We observed in both experimental communities that modifier density had a negative effect on the space clearance rate of the predator. The ability to identify the processes impacted by the modifier allows for a greater understanding of the biology underlying the observed phenomenological change in consumption. In both communities, models that only include pairwise interactions (i.e., predation and competition) were worse at describing the observed changes in population densities than models that included any TIM. Furthermore, simulations of community dynamics using pairwise interaction models produced quantitatively different predictions regarding species persistence and dynamics than models that included TIMs. Thus, our study highlights the need to extend past pairs of focal species to understand trophic interactions and provides an experimental approach to quantify the strength and sign of TIMs on predator foraging.

### Trophic interaction modifications decrease predator consumption

In both experimental communities, the top four best-fitting models all included a modifier density-dependent scaling parameter for space clearance rate (Table 2). This provides substantial support that the modifier is indeed impacting the predator’s space clearance rate. This study focused on a single predator-modifier species combination, however, the alteration of the space clearance rate observed in our communities may be a common effect of modifier species. Kratina *et al*. (2007) demonstrated that in protist communities a predator’s functional response was significantly different in the presence and absence of a non-prey species in a manner consistent with a decrease in the space clearance rate.

Furthermore, using time series data van Veen *et al*. (2005) observed that the dynamics of a host-parasitoid community with a modifier species were better explained by a model that included a TIM on space clearance rate than a pairwise model. Although the top-performing models in this study did not include a TIM on handling time, it remains a possibility with other predator-modifier combinations. For example, in plants and anurans, co-occurring species can alter morphological and chemical traits of the focal species and change its susceptibility to herbivory/predation (Relyea 2002; Castagneyrol *et al*. 2017; Mutz *et al*. 2022). In these systems, the presence of a third species impacts prey traits related to palatability, suggestive of a modified handling time (Box 1). However, to our knowledge, aside from multispecies functional response studies (e.g., Smout & Lindstrøm 2007), an effect of a modifier on handling time has yet to be quantified.

Modifier density had a negative effect on predator space clearance rate and a concomitant negative effect on consumption. The space clearance rate is influenced by the traits of the predator and prey (Box 1) and the environmental matrix (Mocq *et al*. 2021), all of which may have been altered by the modifier. Previous studies have observed that increasing structurally environmental complexity decreases a predator’s space clearance rate (Manatunge *et al*. 2000; Hauzy *et al*. 2010). The presence of the modifier species may have introduced a similar type of complexity making it easier for the prey to evade the predator (e.g., hiding amongst the modifier). Alternatively, a modifier species may directly interfere with the signals (e.g., semiochemicals) a predator uses to find its prey (Vos *et al*. 2001; Hamels *et al*. 2004; Kratina *et al*. 2007). While the design of this study does not allow for the identification of the exact mechanisms of the interaction modification, by identifying the relevant process impacted by the modifier we provide clear targets for future investigations.

The magnitude of the TIM differed between the two prey species. The modifier had a much stronger effect on predator consumption of *Colpidium* than *Dexiostoma* (Figure 2a, c). However, the reason underlying this difference remains unclear. *Paramecium* tends to be more morphologically similar to *Colpidium* than *Dexiostoma* (Appendix Figure S2-1), which may have increased predator interaction with the modifier species (*Paramecium*) or created a cryptic background. However, there are other unmeasured phenotypic traits that may also contribute to the negative effect of our modifier on consumption. An exciting area of future research could be how similarities between modifier and prey species relate to the magnitude and sign of trophic interactions and what traits define these similarities.

### Incorporating trophic interaction modifications to better understand community dynamics

Increased modifier density weakened trophic interaction strength (i.e., reduced consumption) in our experimental communities. As communities that contain fewer strong interactions tend to be more stable (McCann *et al*. 1998), our observation that a modifier species can reduce interaction strength suggests their presence may stabilize community dynamics. This stabilizing effect has widespread theoretical support (Bairey *et al*. 2016; Grilli *et al*. 2017; Singh & Baruah 2021; Gibbs *et al*. 2023) . In this study we do not see stabilization of the dynamics due to the strong competitive interaction between the modifier and the prey species, however such an effect has been observed in a previous study (van Veen *et al*. 2005). The impact of the modifier on space clearance rate suggests that TIMs may be particularly important in communities where predation pressure pushes prey populations to very low densities. In such cases, the presence of the modifier may create a refuge that prevents prey extinction, which may allow for prolonged predator persistence. Predator refugia in a multispecies context have previously been observed with generalist predators that “switch” between prey species depending on abundance (Murdoch & Oaten 1975). Our results suggest that refugia may also exist for the prey of specialized consumers when other species are present that interfere with the interaction. This study utilizes small community motifs, to study the role of TIMs on communities. Increasing experimental evidence indicates that quantifying the interactions between a trio of species allows for the accurate prediction of complex microbial communities dynamics (Friedman *et al*. 2017; Skwara *et al*. 2023; Ishizawa *et al*. 2024). As such, we believe the TIMs observed in our three species motifs, will be informative of the dynamics in more species-rich communities.

The inclusion of a TIM affected important aspects of the dynamics such as time to extinction and population cycle period, thus only considering pairwise interactions may limit the predictability of community dynamics. Despite a large body of work documenting that TIMs occur frequently and in a diversity of systems (Werner & Peacor 2003), they are rarely accounted for in in species interaction models (Bolker *et al*. 2003; Terry *et al*. 2017). The exclusion of TIMs likely hinders our understanding of many ecosystems and may result in unanticipated changes in population density and community composition (Fox 2023), known as ecological surprises, even in well-studied systems (Doak *et al*. 2008). The inclusion of TIMs could play an important role in managing natural systems as a focus on pairwise interactions neglects the stabilizing role of species not directly involved in a trophic interaction (Kadoya *et al*. 2018; Pearson *et al*. 2022). In such circumstances, the stabilizing species may not be deemed to be of management value and allowed to be overexploited eventually resulting in the unexpected, and likely, unexplained collapse of a species of value (Estes *et al*. 1998). Our results suggest that expanding the species interactions framework past pairwise interactions is needed to accurately predict community dynamics and has the potential to aid in understanding previously unexplained ecological patterns (Bolker *et al*. 2003).

### Methodological Advances

In this study, we used the response surface approach proposed by (Terry *et al*. 2017) to quantify the simultaneous change in a prey, predator, and modifier across of range of initial densities of all species. Data collection for response surfaces is laborious and simultaneously fitting the data requires complex analytical methods (Okuyama & Bolker 2012), however this approach , has two major advantages. First, using gradients of at least five initial conditions, allows for interpolation of ecological conditions not measured (Collins *et al*. 2022). Second, simultaneous fitting generates more accurate and precise parameter estimates than fitting conditions independently (Uszko *et al*. 2020). While response surfaces have successfully been used to investigate the effect of prey body size (McCoy & Bolker 2008; McCoy *et al*. 2011) and temperature (Gilioli *et al*. 2005; Uszko *et al*. 2020) on predator’s functional response, our study is the first to investigate TIMs. We alleviated the logistic burden of data collection using computer vision tools (Pennekamp & Schtickzelle 2013) enabling our study to include more total observations and a greater number of initial prey densities and environmental conditions. Our results highlight the potential of response surfaces to understand species interactions and predict community dynamics.

Our approach represents an important advancement in the quantification of TIMs as it expands previous efforts which solely documented the presence and magnitude of a modification (Bolker *et al*. 2003; Terry *et al*. 2017). Using this method, we were able to confidently estimate the parameter values for all functional response formulations, which was previously not possible for similarly complex dynamic models (Fox 2023). These parameters not only enable the prediction of community dynamics, but also provide “anchors” to help ground and evaluate theoretical investigations (Levine *et al*. 2017). A second major advancement of this approach is by comparing functional response formulations with different modifier density dependent parameters, we can identify the consumption process impacted by the modifier species. Past experimental efforts have demonstrated that community dynamics are indeed better described by a model that includes a TIM, however they did not rigorously test different possible biological processes (van Veen *et al*. 2005). Although the nature of the models prevents the identification of the exact modifier induced change, our approach narrows the possible mechanisms to those related to space clearance rate and allows for more directed follow-up experiments. Since its derivation by Holling, the functional response approach utilized in this study has been a favourite instrument in the community ecologist’s toolbox (Jeschke *et al*. 2002; DeLong 2021). We believe that our findings demonstrate that despite many years of research about the functional response, there remains much to be learned, especially when considering the wider food web context (Abrams 2022).

## Conclusions

Our study clearly demonstrates the importance of TIMs in understand and predict community dynamics. Additionally, we illustrate the effectiveness of a response surface approach to identify and quantify these interactions. The simulation component of our study suggests that TIM can affect community stability and thus may represent an important component to understanding how species coexist. Empirical studies are now needed to investigate how the short-term TIMs observed in this study manifest over longer time scales. This information can then be used to validate and adjust current theoretical models.

TIMs are likely widespread in multi-species and multi-trophic ecosystems (Werner & Peacor 2003; Golubski & Abrams 2011). However, not all modifications will be of equal magnitude and while some interactions will have significant impacts, other modifications will have a negligible effect on the strength of a trophic interaction. An important next step is to understand when TIMs will significantly influence trophic interactions and community dynamics, and when they can be safely ignored. In a rapidly changing world, it is increasingly urgent that we understand the ecosystems we aim to manage to avoid adverse ecological surprises. TIMs represent an important, but often unobserved element linking species, and in some systems may be key to understanding and forecasting dynamics.

## Acknowledgments

This work was supported by the Swiss National Science Foundation (grant 310030_197811 awarded to FP). We would like to thank Y. Choffat for laboratory assistance throughout the study. We also thank the members of the Biotic Responses to Environmental Change and Predictive Ecology groups at the University of Zurich for constructive comments on the manuscript.

## Text Boxes

### Box 1 Functional response parameters and their biological interpretation

A functional response describes the change in predator consumption rate with prey density. This model has been extensively used to empirically quantify the interaction strength between species pairs of interest and investigate how biotic (Vucic-Pestic *et al*. 2011; Baudrot *et al*. 2016; Stouffer & Novak 2021) and abiotic factors (Rall *et al*. 2012; Kreuzinger-Janik *et al*. 2019) alter trophic interactions. Functional responses range from linear to non-linear, the latter providing greater biological realism (but see Coblentz *et al*. 2022). Furthermore, they provide estimates of trophic interaction parameters that can easily be incorporated into dynamic models, unlike other measures of interaction strength (i.e., Paine’s index, Berlow *et al*. 2004).

The basic non-linear functional response model is a hyperbolic saturating curve described by Holling (1959) as a Type II response:

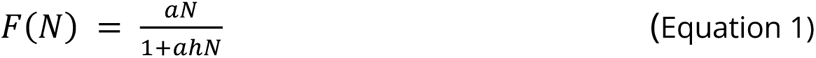

In this formulation, the consumption of the predator (F(N)) is modelled by a function of the density of the prey species (N), and two additional parameters, handling time (h) and space clearance rate (a) of the predator (Figure 1b).

The time required to process a prey individual is called handling time. As handling time increases, fewer prey can be processed, and the maximum ingestion rate of the predator will decrease (Figure 1b-i). The handling time is determined by processes such as the time the predator takes to kill, ingest, and digest prey as well as the time to recover from the consumption and the time spent on unsuccessful attacks. Handling time is influenced by traits and behaviours of both the predator and prey, such as the prey expression of protective defences (Ramos-Jiliberto *et al*. 2008) or predator alterations of gut retention time (Jumars 2000). Interference from conspecific predators can also impact handling time (Beddington 1975; DeAngelis *et al*. 1975; Crowley & Martin 1989). Alterations of this parameter will influence the peak density of prey and predator populations due to the strong relationship between handling time and maximum ingestion rate. If increases in handling time are large, it may prevent the predator from consuming enough resources to meet its energetic demands, potentially leading to predator extinction (Vos *et al*. 2001).

The space clearance rate describes the area (or volume) in which all prey individuals are ‘cleared’ over a set time interval (DeLong 2021) and primarily determines the rate of consumption at low prey densities (Figure 1b-ii). This parameter is influenced by how both predator and prey move in space, the distance from which the predator can detect the prey, the probability of attack, and the probability of a successful attack. The traits and behaviours of both species will impact the space clearance rate, such as the prey altering their movement to evade the predator (Langer *et al*. 2019). Changes in the environmental matrix, such as increased habitat complexity (Barrios-O’Neill *et al*. 2015), can also affect the space clearance rate. Alterations in this parameter will have the greatest effect on consumption at densities where prey are most vulnerable to extinction. As such, a reduction in the space clearance rate could help to stabilize the community dynamics whereas an increase could jeopardize the persistence of the prey species.

A limitation of Type II functional response models is the assumption that space clearance is independent of prey density. Empirical evidence supports density-dependent rates in some conditions (Kalinkat *et al*. 2023). For example, predators may switch from passive to active feeding as prey becomes more abundant (Kiorboe *et al*. 2018). Although the functional response derived by Real (1977, 1979) is often used to model prey density-dependent space clearance rates, we use an alternative form which suppresses the space clearance rate at low prey densities and converges on a type II response at high densities (Hassell *et al*. 1977; DeLong 2021).

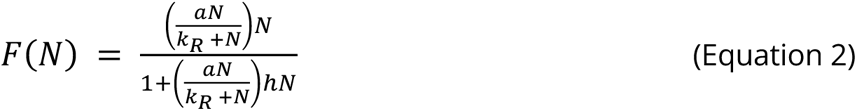

In this model form (hereafter referred to as a generalized functional response) N, a, and h have the same definition as the Type II model. The additional k_R_ parameter is the prey density where the realized space clearance rate reaches half of the maximum. This form prevents biologically unrealistic parameter estimates by constraining the space clearance rate to a maximum value (DeLong 2021), unlike the functional response formulation of Real (1979). Density-dependent space clearance rates produce sigmoidal functional responses, with low consumption at low resource densities. Such a response is generally considered to produce more stable community dynamics than when the space clearance rate is constant (Uszko *et al*. 2015).

## Notes

https://github.com/k-lemmen/Trophic-interaction-modification

### Competing Interest Statement

The authors have declared no competing interest.

## References

Abrams, P.A. (1995). Implications of Dynamically Variable Traits for Identifying, Classifying, and Measuring Direct and Indirect Effects in Ecological Communities. Am. Nat., 146, 112–134.

Abrams, P.A. (2022). Food web functional responses. Frontiers in Ecology and Evolution, 10.

Aho, K., Derryberry, D. & Peterson, T. (2014). Model selection for ecologists: the worldviews of AIC and BIC. Ecology.

Arditi, R. & Ginzburg, L. (2012). How Species Interact: Altering the Standard View on Trophic Ecology. Oxford University Press.

Bairey, E., Kelsic, E.D. & Kishony, R. (2016). High-order species interactions shape ecosystem diversity. Nat. Commun., 7, 12285.

Barbosa, P., Hines, J., Kaplan, I., Martinson, H., Szczepaniec, A. & Szendrei, Z. (2009). Associational Resistance and Associational Susceptibility: Having Right or Wrong Neighbors. Annu. Rev. Ecol. Evol. Syst., 40, 1–20.

Barrios-O’Neill, D., Dick, J.T.A., Emmerson, M.C., Ricciardi, A. & MacIsaac, H.J. (2015). Predator-free space, functional responses and biological invasions. Funct. Ecol., 29, 377–384.

Baudrot, V., Perasso, A., Fritsch, C., Giraudoux, P. & Raoul, F. (2016). The adaptation of generalist predators’ diet in a multi-prey context: insights from new functional responses. Ecology, 97, 1832–1841.

Beddington, J.R. (1975). Mutual Interference Between Parasites or Predators and its Effect on Searching Effciency. J. Anim. Ecol., 44, 331–340.

Berlow, E.L., Neutel, A.-M., Cohen, J.E., De Ruiter, P.C., Ebenman, B., Emmerson, M., et al. (2004). Interaction strengths in food webs: issues and opportunities. J. Anim. Ecol., 73, 585–598.

Billick, I. & Case, T.J. (1994). Higher Order Interactions in Ecological Communities: What Are They and How Can They be Detected? Ecology, 75, 1530–1543.

Bolker, B., Holyoak, M., Křivan, V., Rowe, L. & Schmitz, O. (2003). Connecting theoretical and empirical studies of trait-mediated interactions. Ecology, 84, 1101–1114.

Bolker B, R Development Core Team (2022). _bbmle: Tools for General Maximum Likelihood Estimation_. R package version 1.0.25, <https://CRAN.R-project.org/package=bbmle>.

Bolker, B.M. (2008). Ecological Models and Data in R. Princeton University Press.

Burnham, K.P. & Anderson, D.R. (2002). Model Selection and Multimodel Inference. Springer New York.

Castagneyrol, B., Bonal, D., Damien, M., Jactel, H., Meredieu, C., Muiruri, E.W., et al. (2017). Bottom-up and top-down effects of tree species diversity on leaf insect herbivory. Ecol. Evol., 7, 3520–3531.

Coblentz, K.E., Novak, M. & DeLong, J.P. (2022). Predator feeding rates may often be unsaturated under typical prey densities. Ecol. Lett.

Collins, S., Whittaker, H. & Thomas, M.K. (2022). The need for unrealistic experiments in global change biology. Curr. Opin. Microbiol., 68, 102151.

Crowley, P.H. & Martin, E.K. (1989). Functional Responses and Interference within and between Year Classes of a Dragonfly Population. J. North Am. Benthol. Soc., 8, 211–221.

Daugaard, U., Petchey, O.L. & Pennekamp, F. (2019). Warming can destabilize predator–prey interactions by shifting the functional response from Type III to Type II.

DeAngelis, D.L., Goldstein, R.A. & O’Neill, R.V. (1975). A Model for Tropic Interaction. Ecology, 56, 881–892.

DeLong, J.P. (2021). Predator Ecology: Evolutionary Ecology of the Functional Response. Oxford University Press.

DeLong, J.P. & Vasseur, D.A. (2012). Size-density scaling in protists and the links between consumer-resource interaction parameters. J. Anim. Ecol., 81, 1193–1201.

Doak, D.F., Estes, J.A., Halpern, B.S., Jacob, U., Lindberg, D.R., Lovvorn, J., et al. (2008). Understanding and predicting ecological dynamics: are major surprises inevitable? Ecology, 89, 952–961.

Estes, J.A., Tinker, M.T., Williams, T.M. & Doak, D.F. (1998). Killer whale predation on sea otters linking oceanic and nearshore ecosystems. Science, 282, 473–476.

Fox, J.W. (2023). The existence and strength of higher order interactions is sensitive to environmental context. Ecology, e4156.

Friedman, J., Higgins, L.M. & Gore, J. (2017). Community structure follows simple assembly rules in microbial microcosms. Nat Ecol Evol, 1, 109.

Gibbs, T., Gellner, G., Levin, S.A., McCann, K.S., Hastings, A. & Levine, J.M. (2023). Can higher-order interactions resolve the species coexistence paradox? bioRxiv.

Gilioli, G., Baumgartner, J. & Vacante, V. (2005). Temperature influences on functional response of Coenosia attenuata (Diptera: Muscidae) individuals. J. Econ. Entomol., 98, 1524–1530.

Golubski, A.J. & Abrams, P.A. (2011). Modifying modifiers: what happens when interspecific interactions interact? J. Anim. Ecol., 80, 1097–1108.

Grilli, J., Barabás, G., Michalska-Smith, M.J. & Allesina, S. (2017). Higher-order interactions stabilize dynamics in competitive network models. Nature, 548, 210–213.

Hamels, I., Mussche, H., Sabbe, K., Muylaert, K. & Vyverman, W. (2004). Evidence for constant and highly specific active food selection by benthic ciliates in mixed diatoms assemblages. Limnol. Oceanogr., 49, 58–68.

Hammill, E., Kratina, P., Vos, M., Petchey, O.L. & Anholt, B.R. (2015). Food web persistence is enhanced by non-trophic interactions. Oecologia, 178, 549–556.

Hassell, M.P., Lawton, J.H. & Beddington, J.R. (1977). Sigmoid Functional Responses by Invertebrate Predators and Parasitoids. J. Anim. Ecol., 46, 249–262.

Hauzy, C., Tully, T., Spataro, T., Paul, G. & Arditi, R. (2010). Spatial heterogeneity and functional response: an experiment in microcosms with varying obstacle densities. Oecologia, 163, 625–636.

Holland, J.N., Ness, J.H., Boyle, A. & Bronstein, J.L. (2005). Mutualisms as consumer–resource interactions. In: Ecology of Predator-Prey Interactions (eds Barbosa P. & Castellanos, I.). Oxford University Press, Oxford, pp. 17–33.

Holling, C.S. (1959). Some characteristics of simple types of predation and parasitism. Can. Entomol., 91, 385–398.

Ishizawa, H., Tashiro, Y., Inoue, D., Ike, M. & Futamata, H. (2024). Learning beyond-pairwise interactions enables the bottom-up prediction of microbial community structure. Proc. Natl. Acad. Sci. U. S. A., 121, e2312396121.

Jeschke, J.M., Kopp, M. & Tollrian, R. (2002). PREDATOR FUNCTIONAL RESPONSES: DISCRIMINATING BETWEEN HANDLING AND DIGESTING PREY. Ecol. Monogr., 72, 95–112.

Jumars, P.A. (2000). Animal Guts as Ideal Chemical Reactors: Maximizing Absorption Rates. Am. Nat., 155, 527–543.

Kadoya, T., Gellner, G. & McCann, K.S. (2018). Potential oscillators and keystone modules in food webs. Ecol. Lett., 21, 1330–1340.

Kalinkat, G., Rall, B.C., Uiterwaal, S.F. & Uszko, W. (2023). Empirical evidence of type III functional responses and why it remains rare. Frontiers in Ecology and Evolution, 11.

Kéfi, S., Berlow, E.L., Wieters, E.A., Navarrete, S.A., Petchey, O.L., Wood, S.A., et al. (2012). More than a meal… integrating non-feeding interactions into food webs. Ecol. Lett., 15, 291–300.

Kehoe, R., Frago, E., Barten, C., Jecker, F., van Veen, F. & Sanders, D. (2016). Nonhost diversity and density reduce the strength of parasitoid-host interactions. Ecol. Evol., 6, 4041–4049.

Keitt, T.H. (2017). odeintr: C++ ODE Solvers Compiled on-Demand. R package version 1.7.1.

Kiorboe, T., Saiz, E., Tiselius, P. & Andersen, K.H. (2018). Adaptive feeding behavior and functional responses in zooplankton. Limnol. Oceanogr., 63, 308–321.

Kratina, P., Vos, M. & Anholt, B.R. (2007). Species diversity modulates predation. Ecology, 88, 1917–1923.

Kreuzinger-Janik, B., Brüchner-Hüttemann, H. & Traunspurger, W. (2019). Effect of prey size and structural complexity on the functional response in a nematode-nematode system. Sci. Rep., 9, 5696.

Langer, S.M., Weiss, L.C., Ekvall, M.T., Bianco, G., Hansson, L.-A. & Tollrian, R. (2019). A three-dimensional perspective of *Daphnia*’s swimming behavior with and without predator cues. Limnol. Oceanogr., 64, 1515–1525.

Letten, A.D. & Stouffer, D.B. (2019). The mechanistic basis for higher-order interactions and non-additivity in competitive communities. Ecol. Lett., 22, 423–436.

Levine, J.M., Bascompte, J., Adler, P.B. & Allesina, S. (2017). Beyond pairwise mechanisms of species coexistence in complex communities. Nature, 546, 56–64.

Manatunge, J., Asaeda, T. & Priyadarshana, T. (2000). The Influence of Structural Complexity on Fish– zooplankton Interactions: A Study Using Artificial Submerged Macrophytes. Environ. Biol. Fishes, 58, 425–438.

Mayfield, M.M. & Stouffer, D.B. (2017). Higher-order interactions capture unexplained complexity in diverse communities. Nat Ecol Evol, 1, 62.

McCann, K., Hastings, A. & Huxel, G.R. (1998). Weak trophic interactions and the balance of nature. Nature, 395, 794–798.

McCoy, M.W. & Bolker, B.M. (2008). Trait-mediated interactions: influence of prey size, density and experience. J. Anim. Ecol., 77, 478–486.

McCoy, M.W., Bolker, B.M., Warkentin, K.M. & Vonesh, J.R. (2011). Predicting predation through prey ontogeny using size-dependent functional response models. Am. Nat., 177, 752–766.

Meyer, D., Dimitriadou, E., Hornik, K., Weingessel, A., Leisch, F., Chang, C. C., & Lin, C. C. (2022). Misc Functions of the Department of Statistics, Probability Theory Group (Formerly: E1071), TU Wien (1.7– 11). e1071. [Computer software]. https://CRAN.R-project.org/package=e1071

Mickalide, H. & Kuehn, S. (2019). Higher-Order Interaction between Species Inhibits Bacterial Invasion of a Phototroph-Predator Microbial Community. Cell Syst, 9, 521–533.e10.

Mocq, J., Soukup, P.R., Näslund, J. & Boukal, D.S. (2021). Disentangling the nonlinear effects of habitat complexity on functional responses. J. Anim. Ecol., 90, 1525–1537.

Murdoch, W.W. & Oaten, A. (1975). Predation and Population Stability. In: Advances in Ecological Research (ed. MacFadyen, A.). Academic Press, pp. 1–131.

Mutz, J., Heiling, J.M., Paniagua-Montoya, M., Halpern, S.L., Inouye, B.D. & Underwood, N. (2022). Some neighbours are better than others: Variation in associational effects among plants in an old field community. J. Ecol., 110, 2118–2131.

Okuyama, T. & Bolker, B.M. (2007). On quantitative measures of indirect interactions. Ecol. Lett., 10, 264– 271.

Okuyama, T. & Bolker, B.M. (2012). Model-based, response-surface approaches to quantifying indirect interactions. In: Trait-Mediated Indirect Interactions: Ecological and Evolutionary Perspectives (ed. T. Ohgushi, O.S.&. R.D.H.). Cambridge University Press, pp. 186–204.

Paine, C.E.T., Deasey, A. & Duthie, A.B. (2018). Towards the general mechanistic prediction of community dynamics. Funct. Ecol., 32, 1681–1692.

Pearson, D.E., Clark, T.J. & Hahn, P.G. (2022). Evaluating unintended consequences of intentional species introductions and eradications for improved conservation management. Conserv. Biol., 36, e13734.

Pennekamp, F., Griffiths, J.I., Fronhofer, E.A., Garnier, A., Seymour, M., Altermatt, F., et al. (2017). Dynamic species classification of microorganisms across time, abiotic and biotic environments-A sliding window approach. PLoS One, 12, e0176682.

Pennekamp, F. & Schtickzelle, N. (2013). Implementing image analysis in laboratory-based experimental systems for ecology and evolution: a hands-on guide. Methods Ecol. Evol., 4, 483–492.

Pennekamp, F., Schtickzelle, N. & Petchey, O.L. (2015). BEMOVI, software for extracting behavior and morphology from videos, illustrated with analyses of microbes. Ecol. Evol., 5, 2584–2595.

Preisser, E.L., Bolnick, D.I. & Benard, M.F. (2005). Scared to death? The effects of intimidation and consumption in predator–prey interactions. Ecology, 86, 501–509.

Rall, B.C., Brose, U., Hartvig, M., Kalinkat, G., Schwarzmüller, F., Vucic-Pestic, O., et al. (2012). Universal temperature and body-mass scaling of feeding rates. Philos. Trans. R. Soc. Lond. B Biol. Sci., 367, 2923–2934.

Ramos-Jiliberto, R., Mena-Lorca, J., Flores, J.D. & Morales-Álvarez, W. (2008). Role of inducible defenses in the stability of a tritrophic system. Ecol. Complex., 5, 183–192.

Real, L.A. (1977). The Kinetics of Functional Response. Am. Nat., 111, 289–300.

Real, L.A. (1979). Ecological Determinants of Functional Response. Ecology, 60, 481–485.

Relyea, R.A. (2002). Competitor-Induced Plasticity in Tadpoles: Consequences, Cues, and Connections to Predator-Induced Plasticity. Ecol. Monogr., 72, 523–540.

Rosenbaum, B. & Rall, B.C. (2018). Fitting functional responses: Direct parameter estimation by simulating differential equations. Methods Ecol. Evol., 9, 2076–2090.

Schmitz, O.J., Beckerman, A.P. & O’Brien, K.M. (1997). Behaviorally mediated trophic cascades: effects of predation risk on food web interactions. Ecology, 1388–1399.

Schmitz, O.J., Krivan, V. & Ovadia, O. (2004). Trophic cascades: the primacy of trait-mediated indirect interactions. Ecol. Lett., 7, 153–163.

Schoener, T.W. & Spiller, D.A. (2012). Perspective: Kinds of trait-mediated indirect effects in ecological communities. A synthesis. In: Trait-Mediated Indirect Interactions: Ecological and Evolutionary Perspectives. Cambridge University Press, pp. 9–27.

Schwarz, G. (1978). Estimating the Dimension of a Model. Ann. Stat., 6, 461–464.

Shen, C., Lemmen, K., Alexander, J. & Pennekamp, F. (2023). Connecting higher-order interactions with ecological stability in experimental aquatic food webs. Ecol. Evol., 13.

Singh, P. & Baruah, G. (2021). Higher order interactions and species coexistence. Theor. Ecol., 14, 71–83.

Skwara, A., Gowda, K., Yousef, M., Diaz-Colunga, J., Raman, A.S., Sanchez, A., et al. (2023). Statistically learning the functional landscape of microbial communities. Nat Ecol Evol, 7, 1823–1833.

Smout, S. & Lindstrøm, U. (2007). Multispecies functional response of the minke whale Balaenoptera acutorostrata based on small-scale foraging studies. Mar. Ecol. Prog. Ser., 341, 277–291.

Stouffer, D.B. & Novak, M. (2021). Hidden layers of density dependence in consumer feeding rates. Ecol. Lett., 24, 520–532.

Terborgh, J.W. (2015). Toward a trophic theory of species diversity. Proc. Natl. Acad. Sci. U. S. A., 112, 11415–11422.

Terry, J.C.D., Morris, R.J. & Bonsall, M.B. (2017). Trophic interaction modifications: an empirical and theoretical framework. Ecol. Lett., 20, 1219–1230.

Uszko, W., Diehl, S., Pitsch, N., Lengfellner, K. & Müller, T. (2015). When is a type III functional response stabilizing? Theory and practice of predicting plankton dynamics under enrichment. Ecology, 96, 3243–3256.

Uszko, W., Diehl, S. & Wickman, J. (2020). Fitting functional response surfaces to data: a best practice guide. Ecosphere, 11, 463.

van Veen, F.J.F., van Holland, P.D. & Godfray, H.C.J. (2005). Stable Coexistence in Insect Communities Due to Density- and Trait-Mediated Indirect Effects. Ecology, 86, 3182–3189.

Vos, M., Berrocal, S.M., Karamaouna, F., Hemerik, L. & Vet, L.E.M. (2001). Plant-mediated indirect effects and the persistence of parasitoid-herbivore communities. Ecol. Lett., 4, 38–45.

Vucic-Pestic, O., Ehnes, R.B., Rall, B.C. & Brose, U. (2011). Warming up the system: higher predator feeding rates but lower energetic efficiencies. Glob. Chang. Biol., 17, 1301–1310.

Wasserman, R.J., Alexander, M.E., Dalu, T., Ellender, B.R., Kaiser, H. & Weyl, O.L.F. (2016). Using functional responses to quantify interaction effects among predators. Funct. Ecol., 30, 1988–1998.

Werner, E.E. & Peacor, S.D. (2003). A Review of Trait-Mediated Indirect Interactions in Ecological Communities. Ecology, 84, 1083–1100.

Werner, E.E. & Peacor, S.D. (2006). Lethal and nonlethal predator effects on an herbivore guild mediated by system productivity. Ecology, 87, 347–361.

Wirsing, A.J., Heithaus, M.R., Brown, J.S., Kotler, B.P. & Schmitz, O.J. (2021). The context dependence of non-consumptive predator effects. Ecol. Lett., 24, 113–129.

Wootton, J.T. (1993). Indirect Effects and Habitat Use in an Intertidal Community: Interaction Chains and Interaction Modifications. Am. Nat., 141, 71–89.

