## Supplemental Materials for "Food web context modifies predator foraging and weakens trophic interaction strength"

### **Appendix S1 – Experimental Methods Supplementary Information**

#### S1a: Ciliate Culture Maintenance

All species were cultured in organic protozoan pellet medium (Carolina Biological Supply Company; concentration of 0.55 g/L, see Altermatt *et al.* 2015) bacterized with *Bacillus subtilis*, *Serratia fonticola,* and *Brevibacillus brevis* both before and during the experiment. The prey and modifier species were maintained in 100mL stock cultures before the experiment. To standardize the condition of the culture during the experiment we ensured the density of the stocks had stabilized at carrying capacity seven days (+/- two days) before the experiment. Predator stock cultures were maintained in 4mL of media in 6-well plates and were fed *Dexiostoma* ad libitum. Before the experiment, predators were moved from the stock cultures to a holding plate with reduced prey densities to ensure the observation of consumption. To do so, three days before the experimental trial block, 224 predators (i.e., the number needed for a block) were manually isolated from the maintenance plates and were haphazardly allocated to one of eight wells on a 12-well plate. We added 2 mL of protist pellet media (Carolina Biological Supply Company; concentration of 0.55 g/L) containing *Dexiostoma* at a density of approximately 3000 individuals/mL, consistent with the conditions of the maintenance plates. Twenty-four hours before the experiment, almost all the prey in the holding plates had been consumed. At this point, we added another 1 mL of *Dexiostoma* to each well so that the density was approximately 1000 individuals/mL in each well. When the prey was collected for the trials some *Dexiostoma* individuals were present in each well suggesting that starvation had not occurred. As such the predators entered the experiment in a consistent physical state that neither represents starvation nor complete satiation.

#### S1b: Functional Response Observation and Block Design

For each community, we factorially combined (Figure S1) the starting densities of the prey (9 levels), modifier (7 densities) and predator (2 levels) to generate a response surface for consumption and competition. To ensure a feasible workload we used a different number of technical replicates for the prey only and prey + modifier trials. As additional species are expected to increase the observed variation, we used six technical replicates for experimental units with all three species and three technical replicates for units with one or two species (Figure S1, fill colour of box/number). Due to the density of stock cultures, it was not logistically possible to combine the highest densities of the modifier with the highest densities of the prey without exceeding the volume of the experimental unit (1 mL). As such some prey-modifier combinations could not be observed for this experiment (*Colpidium* community = 2 combinations, *Dexiostoma* community = 4 combinations, Figure S1-1).

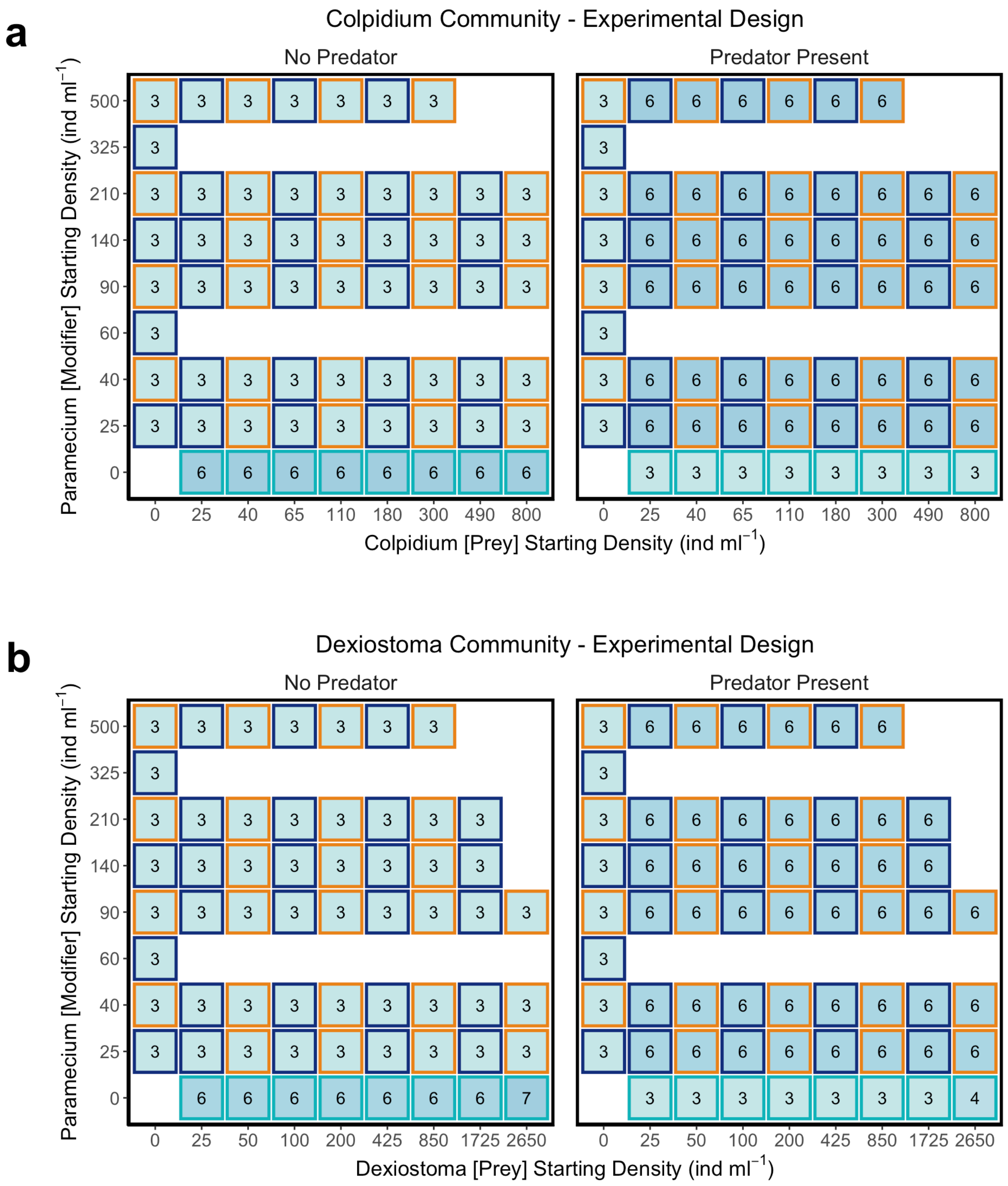


Figure S1-1 – Visual representation of the functional response surface experimental design for a) the *Colpidium* community and b) *Dexiostoma* community. For each experiment, experimental units were observed without a predator (left big box) to estimate growth as well as intra- and interspecific competition parameters, and with a predator (right big box) to estimate functional parameters. Each coloured box represents a combination of prey and modifier densities. The number inside the box indicates the number of technical replicates (light blue <4, blue >4) and the border colour represents the block to which the prey-modifier combination belongs (teal = block 1, dark blue = block 2, orange = block 3).

It was not possible to conduct all experimental units at once, as such we used a block design to collect observations. For both communities, we used three blocks: (i) observations of prey with and without the predator (absence of the modifier), (ii) observations of half the prey and modifier combinations with and without the predator and (iiI) observations of the other half the prey levels and modifier combinations with and without the predator. The observations of the first block (prey only) were collected approximately 2-4 weeks before the other blocks as part of a different experiment. We divided the second and third blocks in a manner that ensured the entire range of prey-modifier combinations was included in each block and that the replicates were spread evenly. One-third of the replicates from the second or third block were run each day over six days alternating between the second and the third block. Prey growth in the absence of the modifier and the predator was measured in all three blocks, which explains the greater number of technical replicates. As most parameters did not significantly differ between periods (Figure S1-2), except save *Dexiostoma* growth rate which was qualitatively similar, we felt comfortable using predator consumption in the absence of the modifier from the earlier experiment.

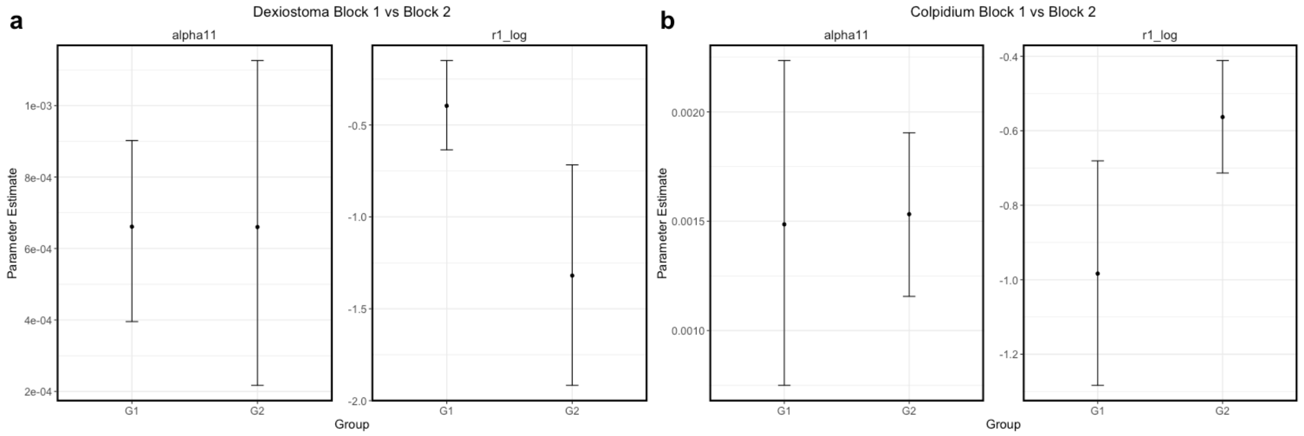


Figure S1-2 – Parameter estimates with 95% confidence intervals simulated using population prediction intervals for the intra-specific competition (alpha11) and growth (r1_log) of a) *Dexiostoma* and b) *Colpidium*. The group label "G1" represents block 1 and "G2" represents block 2/3. Note that estimates for growth are on the log scale.

S1c: Estimating Prey and Modifier Density Using Video Analysis

For each sampling event, the content of the well was gently agitated using a pipette, a subsample of 680μL was then mounted onto a glass slide and covered with a glass lid. We took three non-overlapping videos (at 25 frames per second) of each slide at 8X magnification on a stereomicroscope (Leica M205 C) mounted with a digital CMOS camera (Hamamatsu Orca Flash 4.0 C11440, Hamamatsu Photonics, Japan) with dark field illumination. As the frame of the video was slightly smaller than the glass side chamber, in total we captured 486μL of media with video setup. We processed the videos using the R package BEMOVI (Pennekamp *et al.* 2015) to estimate species abundance. We used the mean of the three videos as our measure of density, if no individuals were detected, we assigned a zero.

For species classification, we trained a support vector machine (SVM) classifier daily with the R package e1071 (Meyer et al., 2022). We generated the training set daily to ensure that if plastic changes in morphology occurred they would not impact classification. We trained the classifier with the videos from that day of experimental units that only contained the prey or the modifier (i.e. growth control or functional response units). Twenty morphological and movement features extracted from an established classification pipeline (Pennekamp *et al.* 2017) were selected to train the SVM to distinguish among classes based on information about body size and movement patterns (Table S1). The classification error in the *Dexiostoma-Paramecium* community was 0.1% and 0.3% respectively. The classification error in the *Colpidium-Paramecium* community was 0.23% and 1% respectively.

Table S1-1: Morphological and movement features selected for use in daily support vector machine classification.

| CODE | Measurement method |
|---|---|
| mean_area | Mean area of particle across trajectory |
| sd_area | Standard deviation of particle area |
| mean_perimeter | Mean length of perimeter of particle |
| sd_perimeter | Standard deviation of particle perimeter length |
| mean_major | Mean length of major axis of ellipse fitted to particle |
| sd_major | Standard deviation of length of major axis |
| mean_ar | Mean aspect ratio of particle |
| sd_ar | Standard deviation of particle aspect ratio |
| mean_turning | Numeric mean of particle direction |
| sd_turning | Circular standard deviation of particle direction |
| gross_disp | Gross displacement (sum of all steps of a trajectory) |
| max_net | Maximum net displacement |
| net_disp | Net displacement |
| net_speed | Net displacement travelled between frames |
| max_step | Maximum step length |
| min_step | Minimum step length |
| sd_step | Standard deviation of step length |
| sd_gross_speed | Standard deviation of distance travelled between frames |
| max_gross_speed | Maximum distance travelled between frames |
| min_gross_speed | Minimum distance travelled between frames |

S1d: Functional Response Parameter Estimation using Numerical Simulations

To estimate the parameter values that best described the observed change in species density we used a combination of numerical simulations of ODE and an iterative maximum likelihood estimator as described in (Rosenbaum & Rall 2018). This approach repeatedly numerically solves the set of equations 3-5 for all combinations of initial prey, modifier, and predator densities with a different set of parameter values. Each parameter set produces a series of predicted values. The likelihood of each set of predicted values is then calculated by assuming that each final observed prey density is log-normally distributed, while modifier and predator density is Poisson distributed around the respective predicted prey, modifier, or predator density. The set of values with the highest likelihood is chosen by the algorithm as the best fit. We fitted the model with some parameters on the log scale to (i) constrain parameter values to be positive, as negative values have no biological meaning and (ii) improve convergence by reducing the magnitude in which the parameters differed.

To estimate the best parameter values for each functional response form, we initiated the numerical simulations using 98 different combinations of starting parameters (i.e., search grid approach). The parameter value combination with the highest loglikelihood was selected to represent that functional response form in model comparison. This approach is our best effort to find a global minimum and avoid the issue of multiple minima that can occur when multiple parameters are being estimated (Bolker 2008). We observed various amounts of convergence of the multiple starting points to the 'best' parameter estimates (mean 55%, min=3%, max=100%).

To ensure differences between models were due solely to differences in the trophic interaction, and not the estimates of prey and modifier growth and intra/interspecific competition, we fixed these parameters values (r1, r2, $\alpha$11, $\alpha$22, $\alpha$12, $\alpha$21) when fitting the ODE system. To do so, we first fit a Lotka-Volterra competition model (equation 1 excluding the consumption component, and equation 2) using the experimental observations without the predator. These parameter values (Appendix S4) were used for all twelve model fits. Additionally, we fixed the mortality (m) of the predator to = 0.1333 individuals/day based on a pre-experimental trial that observed mortality rates of well-fed predators in the absence of food over 24 hours. Our observed value is within the range of protist mortality rates from the literature (DeLong *et al.* 2015). We also fixed the conversion rate efficiency of the predator (c) for all models to 3.06 e-07 (unitless) based on estimates from pre-experimental trial fits. By fixing some model parameters we reduced the parameters that need to be estimated which helps with the fitting of the model.

S1e: Assessment of community stability

We assessed community stability using different metrics depending on whether (i) at least one or (ii) no species went extinct during the simulated dynamics. If at least one species went extinct, as with the *Colpidium-Paramecium-Spathidium* community, we assessed the time to extinction for each species. Populations were considered extinct when population densities were less than 0.1 individuals/mL. The 95% confidence intervals were extracted from population prediction intervals (Bolker 2008). If all species persisted throughout the simulated dynamics, as with the *Dexiostoma-Paramecium-Spathidium* community, we assessed the period and the amplitude of oscillations. To quantify the oscillation period we calculated the number of days between cycle peaks for the prey species (similar periods are observed for all species). Oscillation amplitude represents the mean amplitude throughout the 1100-day dynamic period. To determine the confidence intervals for period and amplitude we simulated the dynamics of the prey population 999 times by randomly drawing model parameter combinations from a multivariate normal distribution (mean = estimated parameter values, variance = covariance matrix of model fit). For each simulated dynamic we calculated the period and amplitude, confidence intervals represent the 2.5 and 97.5 quartiles of the 999 values.

S1f: Sensitivity analysis of conversion efficiency on simulated community dynamics

For the simulated dynamics of both communities, the conversion efficiency of the predator (i.e. the number of new cells produced for each prey cell consumed) was fixed at a value of 0.007. To understand the impact of predator conversion efficiency on the observed dynamics we performed a sensitivity analysis using a range of values (0.005-0.05) observed in protist predators (DeLong & Vasseur 2012). For both communities, we simulated the dynamics including the confidence interval for the best overall and best pairwise model for 100 different values of the conversion efficiency. For the *Colpidium-Paramecium-Spathidium* community, we calculated the time to prey extinction and confidence intervals, and for the *Dexiostoma-Paramecium-Spathidium* community, we calculated point estimate and confidence intervals of the period and amplitude for the prey population for the best overall and best pairwise model for each conversion efficiency value. Time to extinction, cycle period and amplitude, as well as the associated confidence intervals, were calculated as described in Appendix S1e.

### Appendix S2 – Supplementary Results

S2a: Modifier-Prey Interaction

We fit a Lotka-Volterra model to the modifier-prey observations without the predator to estimate the fixed parameter values for the functional response fitting. In the *Colpidium-Paramecium* pair, the growth rate of *Colpidium* was more than double *Paramecium* (Parameter estimates in Table S2). The per capita effect of *Paramecium* on *Colpidium* growth rate was stronger than *Colpidium* on itself. In contrast, the per capita effect of *Colpidium* on *Paramecium* was less than half of the effect *Paramecium* had on itself. In the *Dexiostoma-Paramecium* pair, *Paramecium* grew very slowly. *Paramecium* had a very strong per capita effect on its growth rate, and there was a slight positive effect of *Dexiostoma* on *Paramecium*. In comparison, *Dexiostoma* had faster growth rates than *Paramecium*, a weak effect on its growth rate and was negatively impacted by the density of *Paramecium*.

Table S2-1: Parameter estimates of the growth rate (r), intraspecific competition coefficient (alpha-ii and alpha-jj) and interspecific competition coefficient (alpha-ij and alpha-ji) between the prey (either *Colpidium* or *Dexiostoma*) and the modifier (*Paramecium*). Parameters are estimated from the competition-only trials (i.e., no predator present) and these values are fixed for the fitting of the ordinary differential equations (Equation 3-5).

| | **Prey** | | | **Modifier** | | |
|---|---|---|---|---|---|---|
| | r | alpha-ii | alpha-ij | r | alpha-jj | alpha-ji |
| *Colpidium-Paramecium* | 3.54E-01 | 1.37E-03 | 1.84E-03 | 1.54E-01 | 1.81E-03 | 8.72E-04 |
| *Dexiostoma-Paramecium* | 4.96E-01 | 4.91E-04 | 2.30E-03 | 2.20E-03 | 2.08E-01 | -3.45E-02 |

### S2b: Parameter values of functional response models

Table S2-2: Parameter estimates of models considered for the *Colpidium-Paramecium-Spathidium* and *Dexiostoma-Paramecium-Spathidium* communities. Functional responses considered may include a modifier effect on space clearance rate (TIM->a) and/or handling time (TIM->h) and modifier inference with the predator (van Veen and Crowley-Martin).

| Prey Species | Model | TIM -> a | TIM -> h | Shape | a | a12 | h | h12 | kr | w |
|---|---|---|---|---|---|---|---|---|---|---|
| *Colpidium* | **2** | **Linear** | | **Type 2** | **0.097** | **-0.00019** | **0.038** | | | |
| | 9 | Linear | | Generalized | 0.106 | -0.00021 | 0.045 | | 0.381 | |
| | 4 | Linear | Linear | Type 2 | 0.097 | -0.00019 | 0.032 | 0.00011 | | |
| | 11 | Linear | Linear | Generalized | 0.106 | -0.00021 | 0.037 | 0.00013 | 0.391 | |
| | 7 | Crowley-Martin | | Type 2 | 0.109 | | 0.029 | | | 0.006 |
| | 5 | van Veen | | Type 2 | 0.116 | | 0.038 | | | 0.007 |
| | 12 | van Veen | | Generalized | 0.125 | | 0.043 | | 0.313 | 0.007 |
| | 6 | van Veen | Linear | Type 2 | 0.112 | | 0.032 | 0.00010 | | 0.006 |
| | 13 | Crowley-Martin | | Generalized | 0.110 | | 0.029 | | 0.004 | 0.006 |
| | 10 | | Linear | Generalized | 0.097 | | 0.016 | 0.00081 | 0.925 | |
| | 3 | | Linear | Type 2 | 0.077 | | 0.008 | 0.00063 | | |
| | 1 | | | Type 2 | 0.064 | | 0.038 | | | |
| | 8 | | | Generalized | 0.070 | | 0.045 | | 0.400 | |
| *Dexiostoma* | **9** | **Linear** | | **Generalized** | **0.281** | **-0.00033** | **0.019** | | **1.73** | |
| | 11 | Linear | Linear | Generalized | 0.295 | -0.00042 | 0.024 | -0.00004 | 1.75 | |
| | 12 | van Veen | | Generalized | 0.287 | | 0.019 | | 1.73 | 0.002 |
| | 13 | Crowley-Martin | | Generalized | 0.270 | | 0.016 | | 1.73 | 0.001 |
| | 8 | | | Generalized | 0.228 | | 0.018 | | 1.92 | |
| | 10 | | Linear | Generalized | 0.237 | | 0.014 | 0.00004 | 2.01 | |
| | 2 | Linear | | Type 2 | 0.214 | -0.00026 | 0.012 | | | |
| | 5 | van Veen | | Type 2 | 0.221 | | 0.012 | | | 0.002 |
| | 4 | Linear | Linear | Type 2 | 0.221 | -0.00030 | 0.016 | -0.00004 | | |
| | 7 | Crowley-Martin | | Type 2 | 0.212 | | 0.010 | | | 0.002 |
| | 6 | van Veen | Linear | Type 2 | 0.230 | | 0.015 | -0.00003 | | 0.003 |
| | 1 | | | Type 2 | 0.172 | | 0.011 | | | |
| | 3 | | Linear | Type 2 | 0.174 | | 0.009 | 0.00002 | | |

S2c: Key morphological and behavioural traits of prey and modifier species

To understand the morphological and behavioural differences between the two prey species (*Colpidium striatum* and *Dexiostoma campylum*) and modifier species (*Paramecium caudatum*) we quantified a subset of traits using monoculture videos collected throughout the experiment (i.e. growth control, and single species functional response treatments). We first calculated the mean of all individuals in each video (*Colpidium*: 216 videos each with approximately 62.7 individuals; *Dexiostoma*: 221 videos each with approximately 190.5 individuals; *Paramecium*: 288 videos each with approximately 59.6 individuals). Trait means are based on video means. The 95% confidence intervals represent nonparametric bootstrapped confidence limits.

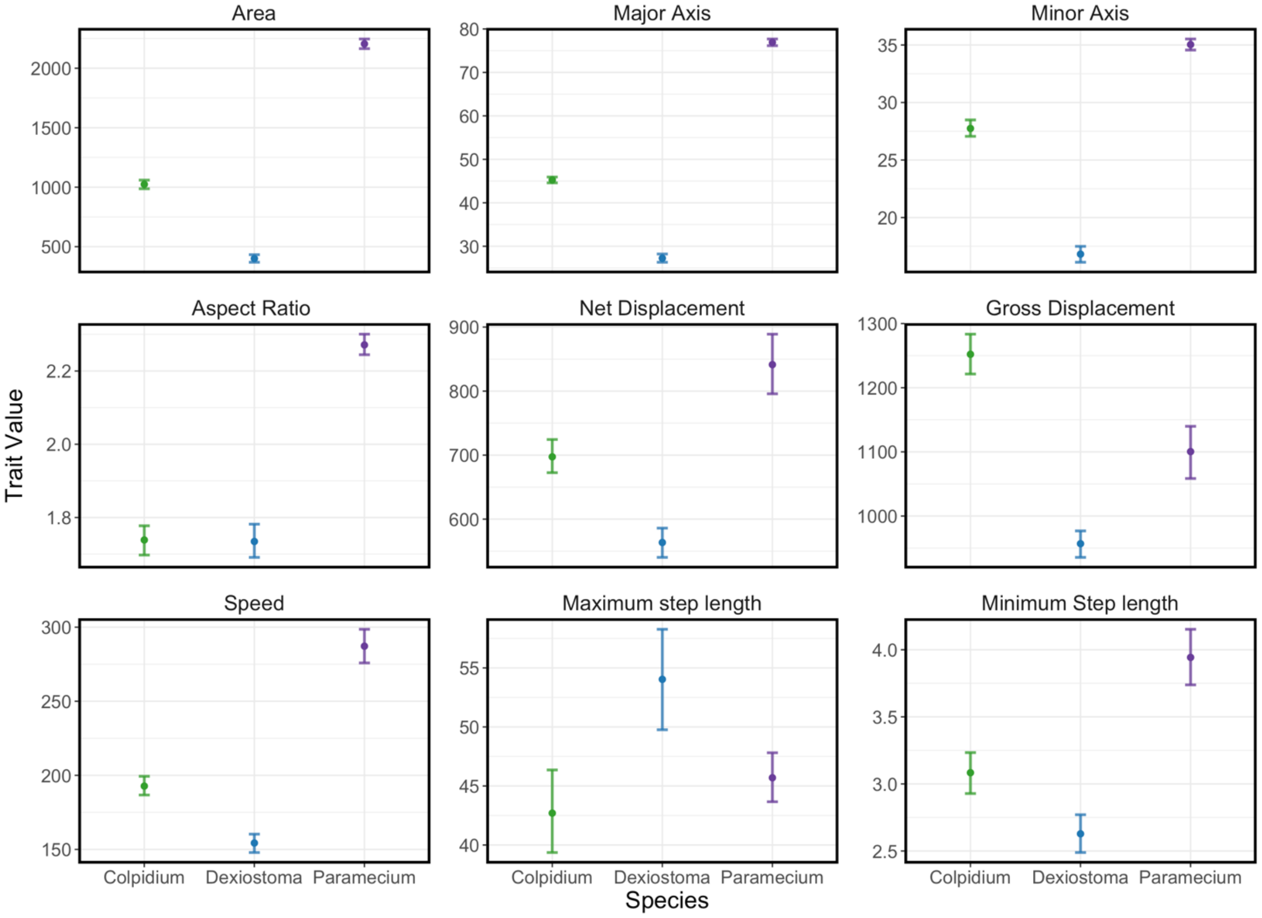


Figure S2-1 – Mean with 95% confidence intervals of morphological and behavioural traits of the two prey species (*Colpidium striatum* and *Dexiostoma campylum*) and modifier species (*Paramecium caudatum*). See Table S1-2 for an explanation of each trait.

### **Appendix S3 – Sensitivity Analysis of Community Dynamics**

For the *Colpidium* community, as conversion efficiencies increase, the difference in the time to extinction between the best overall and best pairwise model increases (Figure S3). We likely see a divergence in time to extinction between models because at larger conversion efficiencies predator population size will increase more rapidly for a single prey item consumed and a larger population will consume more prey individuals. The capacity of the modifier to suppress prey consumption prevents the rapid increase in predator population sizes that occur at higher conversion efficiencies and thus prevents a dramatic rapid decline in the prey population which leads to earlier extinction times.

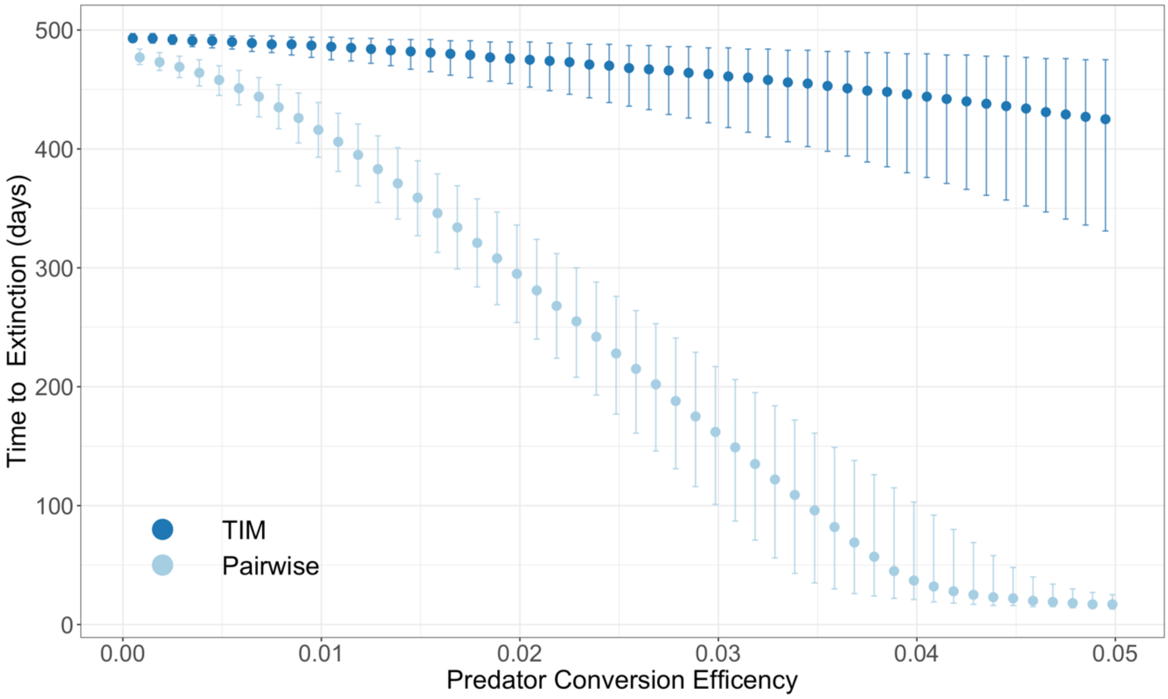


Figure S2-2 – *Colpidium* sensitivity analysis investigating the effect of predator conversion efficiency on the population dynamics regarding time to extinction, for a model that accounts for a trophic interaction modification (TIMs) and a model that only includes pairwise interactions. The 95% confidence intervals were extracted from population prediction intervals.

For the *Dexiostoma* community, as predator conversion efficiencies increase, both the period and the amplitude of the prey population oscillations become smaller (Figure S4). Differences between the overall best model and the best pairwise model are greatest at low predator conversion efficiencies (period =0.004; amplitude = 0.0035), although never significantly different. As predator conversion efficiencies increase the difference between the community dynamics between the trophic interaction modification model and pairwise only model diminishes.

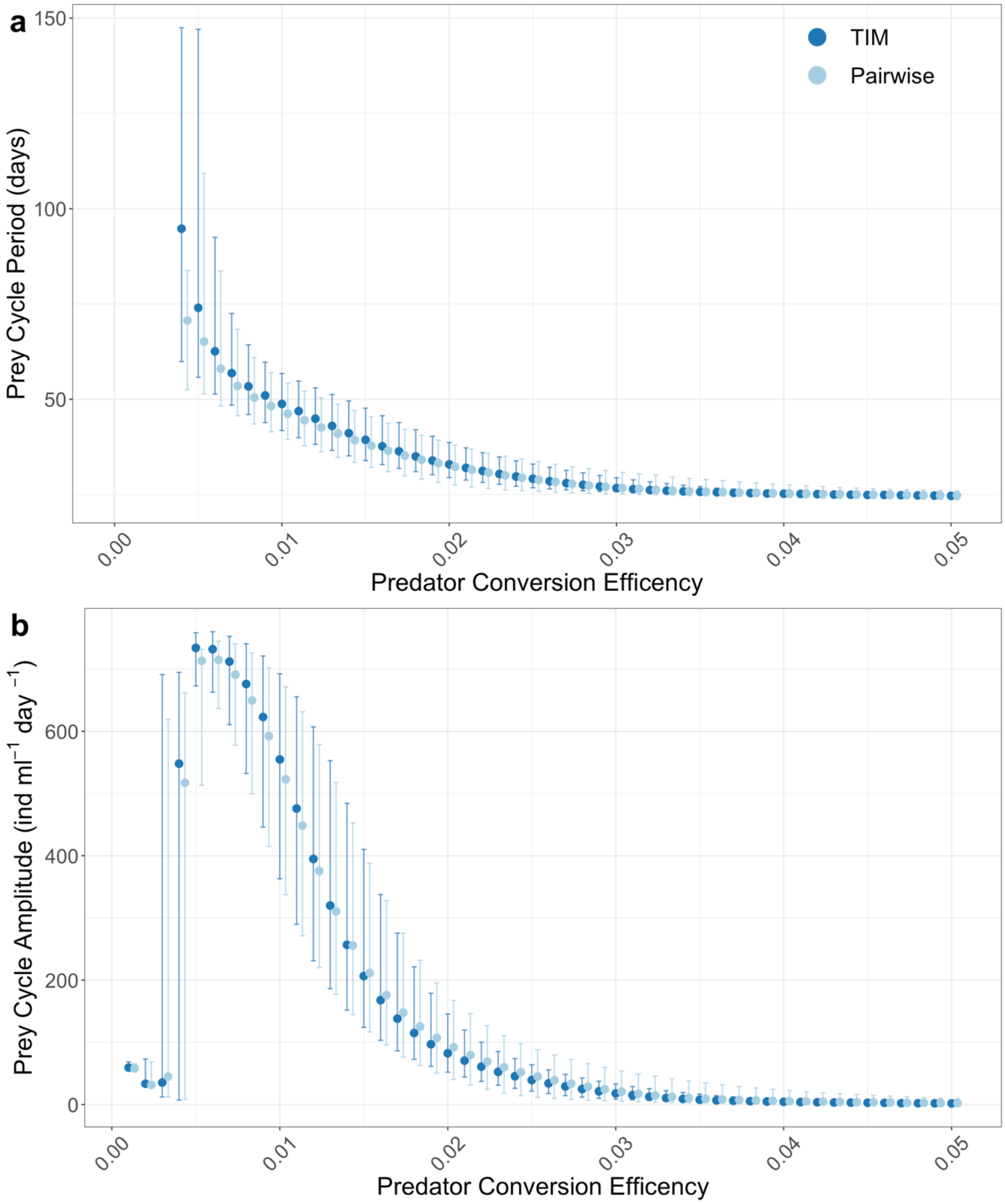


Figure S2-3 – *Dexiostoma* sensitivity analysis investigating the effect of predator conversion efficiency on population dynamics regarding cycle a) Period and b) Amplitude, for a model that accounts for a trophic interaction modification (TIMs) and a model that only includes pairwise interactions. Confidence intervals represent the 2.5 and 97.5 quartiles of 999 simulated dynamics at a given predator conversion efficiency with model parameter combinations drawn from a multivariate normal distribution (mean = estimated parameter values, variance = covariance matrix of model fit).

### Literature Cited


Altermatt, F., Fronhofer, E.A., Garnier, A., Giometto, A., Hammes, F., Klecka, J., *et al.* (2015). Big answers from small worlds: a user's guide for protist microcosms as a model system in ecology and evolution. *Methods Ecol. Evol.*, 6, 218–231.

Bolker, B.M. (2008). *Ecological Models and Data in R*. Princeton University Press.

DeLong, J.P., Gilbert, B., Shurin, J.B., Savage, V.M., Barton, B.T., Clements, C.F., *et al.* (2015). The body size dependence of trophic cascades. *Am. Nat.*, 185, 354–366.

DeLong, J.P. & Vasseur, D.A. (2012). Size-density scaling in protists and the links between consumer-resource interaction parameters. *J. Anim. Ecol.*, 81, 1193–1201.

Meyer, D., Dimitriadou, E., Hornik, K., Weingessel, A., Leisch, F., Chang, C. C., & Lin, C. C. (2022). Misc Functions of the Department of Statistics, Probability Theory Group (Formerly: E1071), TU Wien (1.7–11). e1071. [Computer software]. https://CRAN.R-project.org/packa ge=e1071

Pennekamp, F., Griffiths, J.I., Fronhofer, E.A., Garnier, A., Seymour, M., Altermatt, F., *et al.* (2017). Dynamic species classification of microorganisms across time, abiotic and biotic environments-A sliding window approach. *PLoS One*, 12, e0176682.

Pennekamp, F., Schtickzelle, N. & Petchey, O.L. (2015). BEMOVI, software for extracting behavior and morphology from videos, illustrated with analyses of microbes. *Ecol. Evol.*, 5, 2584–2595.

Rosenbaum, B. & Rall, B.C. (2018). Fitting functional responses: Direct parameter estimation by simulating differential equations. *Methods Ecol. Evol.*, 9, 2076–2090.
